# Possible involvement of ghost introgressions in the striking diversity of Vomeronasal type 1 receptor genes in East African cichlids

**DOI:** 10.1101/2024.12.19.629293

**Authors:** Shunsuke Taki, Zicong Zhang, Mitsuto Aibara, Tatsuki Nagasawa, Masato Nikaido

**Author notes:** Correspondence: Masato Nikaido.

## Abstract

Cichlids that have undergone adaptive radiation are genetically close but exhibit extreme ecological and morphological diversity, making them useful for understanding speciation mechanisms. Vomeronasal type 1 receptors (V1R) are highly conserved among teleost fish at the amino acid sequence level and believed to play a fundamental role in reproduction. We previously reported the surprisingly high sequence diversity of V1Rs among certain cichlid species, suggesting a possible role for V1Rs in their speciation. In this study, we investigated the process of evolutionary diversification of all 6 V1Rs (V1R1-6) by using the genome data of 528 cichlid species, encompassing nearly all lineages. In the case of V1R2, two highly divergent alleles (1.17%: variant sites/CDS length) without recombination were preserved and shared among cichlids found in all of the East African Great Lakes. In the case of V1R6, numerous highly variable alleles that could be derived from multiple recombination events between two highly divergent alleles (1.39%: variant sites/CDS length) were found among the Lake Victoria cichlids. Additionally, we identified highly divergent alleles of V1R1 within the tribe Tropheini, and of both V1R3 and V1R6 within Trematocarini and Ectodini. Because one of the two divergent alleles of these V1Rs emerged rapidly during cichlid evolution, they are likely to have been derived from introgression. However, despite extensive investigations, we could not identify the source lineages for these introgressions, implying that they may have become extinct. This study revealed the potential role of introgression in explaining the remarkable diversity of V1Rs in East African cichlids.

## Introduction

Cichlids (family Cichlidae) are tropical freshwater fish that inhabit various regions spanning Central and South America to Africa, Madagascar, and India. The total number of cichlid species is estimated to be approximately 3,000, representing approximately 10% of all teleost fish (Kocher, 2004). Notably, approximately half of the extant cichlid species inhabit the East African Great Lakes, specifically Lake Tanganyika, Lake Malawi, and Lake Victoria (Salzburger et al., 2014), where they have emerged through large-scale adaptive radiation in each lake (Kocher, 2004; Salzburger, 2018). These cichlid species are widely used as model organisms for investigating speciation mechanisms, owing to their significant ecological and morphological diversity despite their genetic closeness. Cichlid species usually exhibit prezygotic isolation, ensuring their reproductive isolation from one another (Rometsch et al., 2020). Speciation is facilitated by sexual selection and assortative mating, particularly in environments where geographic isolation is less probable, such as lakes (Seehausen & Wagner, 2014).

Cichlids rely on various sensory systems for their reproductive behavior (Escobar-Camacho & Carleton, 2015). Because cichlids exhibit multiple body colors, their visual system is a subject of ongoing research. Previous studies have shown that the visual system has diversified according to ecological conditions and habitats, and that its environmental adaptation have contributed to speciation (Carleton & Yourick, 2020; Seehausen et al., 2008; Terai et al., 2006). The promotion of speciation through environmental adaptation of sensory systems, termed “sensory drive,” plays a significant role in the speciation of cichlids. For instance, a shift in the peak sensitivity of long-wavelength-sensitive (LWS) opsin in response to the light environment (blue or red light), coupled with a divergence in female preference for male nuptial coloration (blue or red), led to cichlid speciation in Lake Victoria (Seehausen et al., 2008).

Cichlids utilize visual and olfactory cues to recognize different species and individuals (Keller-Costa et al., 2015; Santos et al., 2023). Behavioral experiments on Lake Malawi cichlids have demonstrated the role of olfaction in conspecific recognition (Plenderleith et al., 2005). Additionally, olfactory imprinting during mouthbrooding was observed in Lake Victoria cichlids (Verzijden & ten Cate, 2007). Moreover, compounds that function as pheromones by influencing male behavior and the female reproductive system were found in the urine of Mozambique tilapia (*Oreochromis mossambicus*) (Ashouri et al., 2023; Keller-Costa et al., 2014). Therefore, cichlids largely depend on olfaction for conspecific recognition; however, in contrast to studies of the visual system, research on which olfactory receptor genes and alleles contribute to species recognition and, hence, assortative mating remains limited.

Four types of olfactory receptor genes—olfactory receptor, trace amine-associated receptor, vomeronasal type 1 receptor (V1R), and V2R—have been identified in teleost fish, all of which are expressed in the olfactory neurons (Calvo-Ochoa et al., 2021). Of these, the V1R gene, also called olfactory receptor related to class A (ORA), is considered to play a role in pheromone detection in fish, as it is homologous to the mammalian V1R, a known pheromone receptor gene (Del Punta et al., 2002; Pfister & Rodriguez, 2005). Although mammalian V1Rs exhibit copy number and sequence diversity, teleost fish V1Rs are highly conserved, with six copies of the gene (V1R1–6) shared across species, and their orthologs are also highly conserved at the amino acid sequence level (Saraiva & Korsching, 2007). Therefore, V1R likely detects hormonal pheromones shared across species, contributing to synchronizing gamete maturation and/or spawning interactions (Ota et al., 2012; Policarpo et al., 2022; Saraiva & Korsching, 2007; Zapilko & Korsching, 2016). The V1Rs of zebrafish have been demonstrated to detect 4-hydroxyphenylacetic acid (4HPAA) and bile acids (Behrens et al., 2014; Cong et al., 2019). Additionally, V1R2- and V1R5-expressing neurons were activated in response to male cichlid urine (Kawamura & Nikaido, 2022). Therefore, V1Rs can significantly contribute to cichlid speciation; however, whether their genetic diversity is associated with speciation or environmental adaptation, as demonstrated in the case of LWS, is unclear.

The V1R genes exhibit sequence-level diversity in East African cichlids (Nikaido et al., 2014). We previously found that certain tribes of Lake Tanganyika cichlids exhibit allelic diversity in the V1R1, V1R3, and V1R6 genes. We also found two alleles of V1R2, and revealed that these alleles are present in cichlid species from different lakes. These alleles differed by >10 nonsynonymous substitutions, suggesting functional divergence. Hence, V1Rs are well-suited for investigating the possibility of olfaction-mediated speciation in cichlids (Santos et al., 2023). However, the limited number of species studied prevents a complete understanding of the origin and development of these divergent alleles.

The recent availability of whole-genome sequencing (WGS) data for >500 cichlid species has enabled detailed comparative genomic analyses across different species (Malinsky et al., 2018; McGee et al., 2020; Ronco et al., 2021). These data enabled us to extensively examine the V1R sequences to elucidate the origin and maintenance of divergent alleles across lineages in a range of morphologically and ecologically diverse cichlids. Accordingly, we conducted a comprehensive analysis of V1R allelic diversity using WGS data from 528 species and 907 specimens of cichlids. We also investigated how sequence-level variations affect the V1R function, particularly its ligand-binding function.

## Materials and Methods

### V1R gene sequences

We used short-read data from 28 tribes, 528 species, and 907 samples of cichlids deposited within the NCBI Sequence Read Archive (Figure S1, Table S1). Low-quality reads were removed using fastp v0.23.2 (Chen et al., 2018), and the filtered reads were mapped to the reference genome (Supplementary Information) using bwa-mem2 v2.2.1 (Vasimuddin et al., 2019) and sorted and indexed using samtools v.1.15 (Danecek et al., 2021). The V1R gene regions were then extracted from the sorted BAM files using samtools v1.15 (Danecek et al., 2021). Variants were called from the extracted BAM files using the HaplotypeCaller tool of GATK v4.3.0 (McKenna et al., 2010). The first round of hard filtering for SNPs and INDELs was performed using the VariantFiltration tool of GATK v4.3.0 (McKenna et al., 2010) based on the following parameters: SNPs (QD<2.0, QUAL<30.0, SOR>4.0, FS>60.0, MQ<40.0, MQRankSum<-12.5, and ReadPosRankSum<-8.0) and INDELs (QD<2.0, QUAL<30.0, FS>200.0, SOR>10.0, and ReadPosRankSum<-20.0). Based on the resulting VCF file, quality recalibration of the BAM files was performed with the BaseRecalibrator and ApplyBQSR tools from GATK v4.3.0 (McKenna et al., 2010). The second round of hard filtering was conducted similarly to the first round. The filtered VCF files were used to generate V1R gene sequences in fasta format using bcftools v1.15 (Danecek et al., 2021). Sequences containing heterozygous variants were phased using HapCUT2 v1.3.3 (Edge et al., 2017). We developed a custom pipeline spanning the process from short-read data download to obtaining V1R gene sequences.

### Phylogenetic analysis

To minimize the computational load of the phylogenetic analysis, we removed duplicate sequences and then randomly extracted 45 sequences from an initial pool of 337 V1R2 sequences, resulting in 5 datasets (Figure 1, Figure S2–6). The datasets were subjected to multiple sequence alignment using MAFFT v7.520 (Katoh & Standley, 2013) with the following options: —maxiterate 16 —localpair. Maximum likelihood trees were constructed using RAxML-NG v1.2.0 (Kozlov et al., 2019) with 1000 bootstrap replicates. The HKY+G4 model was selected using Modeltest-NG v0.1.7 (Darriba et al., 2020) for phylogenetic tree construction. Datasets 1 through 5 exhibited similar topological patterns (Figure S2–6); thus, we only used Dataset 1 (Figure 1). For V1R1 of Tropheini, we used 43 nonduplicated V1R1 sequences and the outgroup *Oreochromis niloticus* V1R1 sequence, totaling 44 sequences (1000 bootstrap replicates, model: HKY+I). For V1R3 of Trematocarini, we used 11 nonduplicated V1R3 sequences and the outgroup *Oreochromis niloticus* V1R1 sequence, totaling 12 sequences (1000 bootstrap replicates, model: HKY+I). For both V1R3 and V1R6, we used 53 nonduplicated sequences (100 bootstrap replicates, model V1R3: HKY+G4, V1R6: GTR+G4). We used the following R packages for visualization: treeio (Wang et al., 2020), ggtree (Yu et al., 2017), readr (Wickham H, Hester J, Bryan J, 2024), dplyr (Wickham H, François R, Henry L, Müller K, Vaughan D, 2023), ggplot2 (Villanueva & Chen, 2019), ggmsa (Zhou et al., 2022), and iTOL v6 (Letunic & Bork, 2024). We calculated the frequencies of each V1R1 and V1R2 allele across cichlid lineages using unphased sequences (V1R1: 907 sequences, V1R2: 907 sequences). We used 1309 sequences to classify the V1R6 haplotypes. Additionally, we used V1R sequences from 102 species of ray-finned fish (Policarpo et al., 2022) and phylogenetic tree data (Chang et al., 2019; Rabosky et al., 2018) to determine the sequence diversity of V1R genes in ray-finned fish. We generated a pairwise similarity matrix for the 102 species using CIAlign v1.0.17 (Tumescheit et al., 2022) and calculated the average similarity. We also calculated pairwise dN/dS ratios using the CODEML program in PAML v4.10.6 (Yang, 2007) and computed the average value.

**Figure 1.**
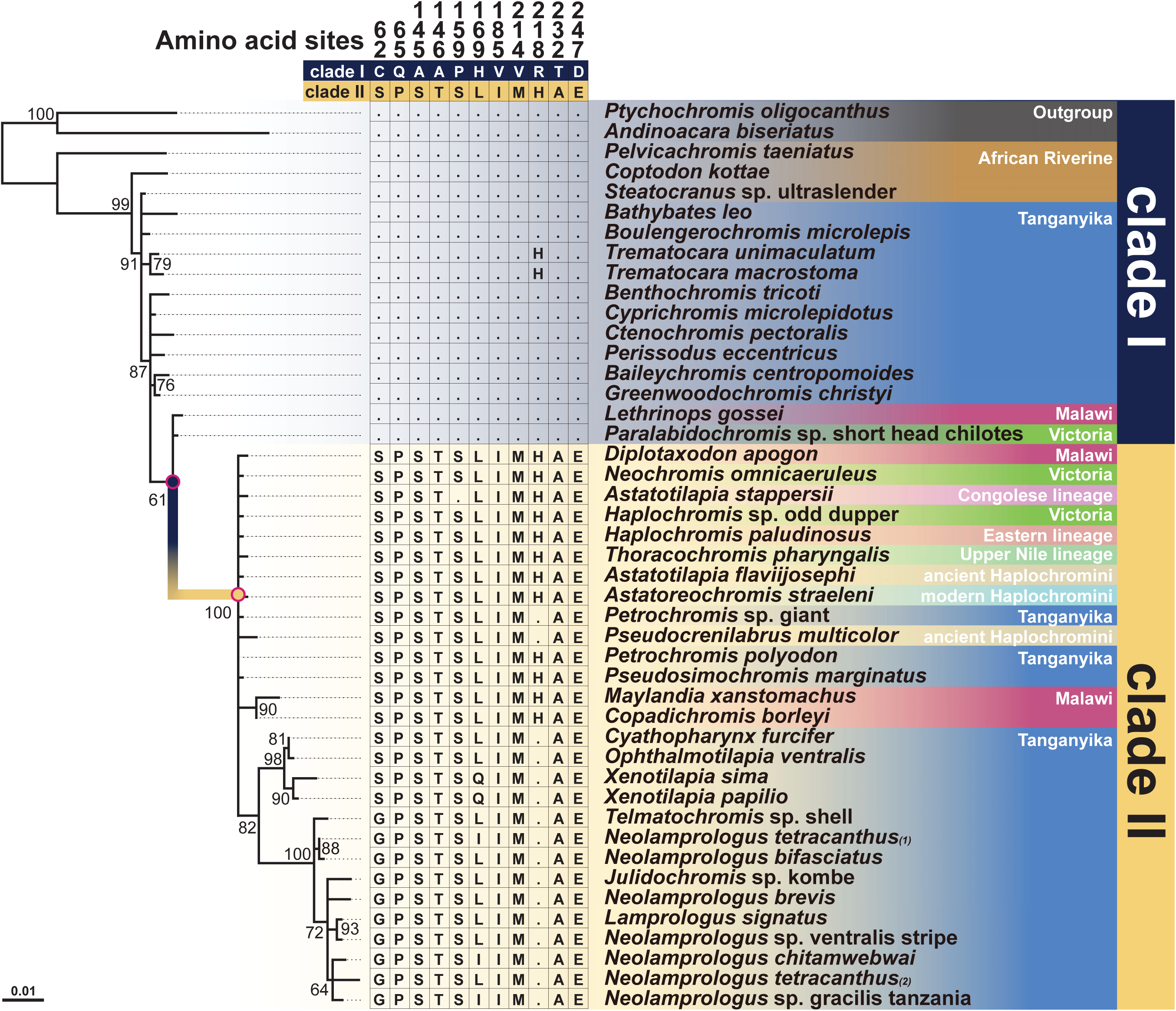
Phylogenetic tree and sequence alignment of the V1R2 gene in cichlids. Bootstrap values are shown only for nodes with support values of ≥60. The sequences are divided into two highly divergent alleles (Clades I and II). Only diagnostic amino acid sites between Clades I and II are shown. Dots indicate identity with the top sequence (Clade I). The phase of each heterozygous allele in the individuals is indicated in parentheses. Observed recombination-derived sequences, which are exceptionally rare, are excluded. Note that sequences from each lake (Tanganyika, Malawi, and Victoria) do not form monophyletic groups.

### Selection analysis

The amino acid sequences translated using Aliview (Larsson, 2014) were subjected to multiple sequence alignment using MAFFT v7.520 (Katoh & Standley, 2013) with the following options: —maxiterate 16 —localpair. The resultant sequence alignment was converted to a codon alignment using PAL2NAL v14 (Suyama et al., 2006). We estimated the dN/dS ratios for branches with an accumulation of nonsynonymous substitutions using the branch model with the codeml program in PAML v4.10.7 (Yang, 2007). We then performed a likelihood ratio test to compare the foreground and background branches and assess the statistical significance of the dN/dS increase in the foreground branch. Clades I and II were established within the V1R2 grouping. The dN/dS ratio was calculated across all five datasets using PAML v4.10.7 (Datasets 1–5, Figure S2–6), and ancestral sequences were reconstructed for the ancestral and descendant nodes of each branch. Analysis of the reconstructed ancestral sequences revealed 11 nonsynonymous substitutions shared across all datasets, which were classified as mutations distinguishing Clade I from Clade II. For V1R1, we defined Clades I and II based on the phylogenetic tree of the V1R1 gene in Tropheini (Figure S9). We identified 11 nonsynonymous substitutions, which were classified as mutations distinguishing Clade I from Clade II.

### Protein modeling and ligand-binding predictions

We modeled the 3D structures of the V1R1, V1R2, and V1R6 proteins using AlphaFold v2.3 (Jumper et al., 2021). We used the most reliable structures to estimate the coordinates of the ligand-binding pockets using Deepsite (Jiménez et al., 2017) in PlayMolecule Viewer (Torrens-Fontanals et al., 2024). We used the obtained coordinates to perform ligand docking simulations in Webina v.1.0.5 (Kochnev et al., 2020) using 4HPAA, a known ligand for zebrafish V1R2 (Behrens et al., 2014), and lithocholic acid, a known ligand for zebrafish V1R1 and V1R6 (Cong et al., 2019). As a result, 5 relative positions (poses) of the ligand in V1R1 (Clade I: 3 poses, Clade II: 2 poses), 18 poses in V1R2 (Clade I: 9 poses, Clade II: 9 poses), and 18 poses in V1R6 (ancestral allele: 9 poses, derived allele: 9 poses) were predicted. Residues located within 5Å of all these poses were designated as ligand-related sites. Furthermore, we predicted transmembrane regions using DeepTMHMM v1.0.39 (Hallgren et al., 2022). The results were visualized using ChimeraX v1.6 (Goddard et al., 2018).

### Recombination rate

We compared the relative recombination rates in the regions surrounding the V1R genes using the recombination rate of *Pundamilia nyerere*i (Feulner et al., 2018) and the genomic position of the V1R genes. We obtained the genomic positions of the V1R genes using BLASTN with the *Pundamilia nyererei* genome assembly (P_nyererei reference v2) (Feulner et al., 2018) as the reference and *Oreochromis niloticus* V1R sequences from NCBI (V1R1, AB671405.1; V1R2, AB670433.1; V1R3, AB704581.1; V1R4, AB704618.1; V1R5, AB704661.1; and V1R6, AB704705.1) as the query. The genomic positions of the V1R1, V1R2, V1R3, V1R4, V1R5, and V1R6 genes were identified as chr13:8171366-8172331, chr13:8175817-8176758, chr13:18527399-18530109, chr13:18531988-18532072, chr2:41585178-41586302, and chr7:27107196-27108122, respectively.

## Results

### Two highly divergent alleles of V1R2 in the cichlids of East African Great Lakes

We constructed a phylogenetic tree for the V1R2 gene using sequence data from 44 species, which were selected from all cichlid lineages (Figure 1). Cichlids in the East African Great Lakes have undergone independent adaptive radiation in each lake, resulting in closely related species in each lake, with species from Lake Malawi and Lake Victoria forming distinct monophyletic groups (Svardal et al., 2021). However, the topology of the V1R2 gene tree did not match the species tree and was divided into Clades I and II with a maximal bootstrap probability (Figure 1). Clade I included cichlids from the outgroup (India and Madagascar), the African Riverine, Lake Tanganyika, Lake Malawi, and Lake Victoria, whereas Clade II included cichlids from Lake Tanganyika, ancient Haplochromini, modern Haplochromini, Lake Malawi, Upper Nile lineage, Eastern lineage, Congolese lineage, and Lake Victoria. Notably both clades contained lineages from all three of the East African Great Lakes. Clades I and II were distinguished based on 11 amino acid substitutions (Figure 1, see Method). Furthermore, the dN/dS ratio of the branch at which Clade II diverged from Clade I was significantly higher than that of the background branches (Table S2), indicating a positive selection acting on the divergent branch.

We calculated the frequencies of the two major alleles (Clades I and II) of V1R2 for each cichlid lineage (Figure 2). In the Lamprologini and Ectodini, which inhabit Lake Tanganyika, V1R2 alleles were fixed in Clade II. In the Haplochromini from Lake Malawi and Lake Victoria, the alleles were not fixed in either clade, resulting in a mix of homozygotes of Clade I or II alleles, along with their corresponding heterozygotes. Despite extensive exploration, Clade II was not detected in the lineages from India, Madagascar, the New World, and the African Riverine. Notably, recombination was not found between the two alleles in all lineages except Cyphotilapiini, despite the alleles coexisting as polymorphisms within populations or species. The polymorphic sites in cichlids that distinguish Clade I from Clade II were highly conserved among noncichlid ray-finned fish, with all V1R2 polymorphic sites being classified into Clade I (Figure S7).

**Figure 2.**
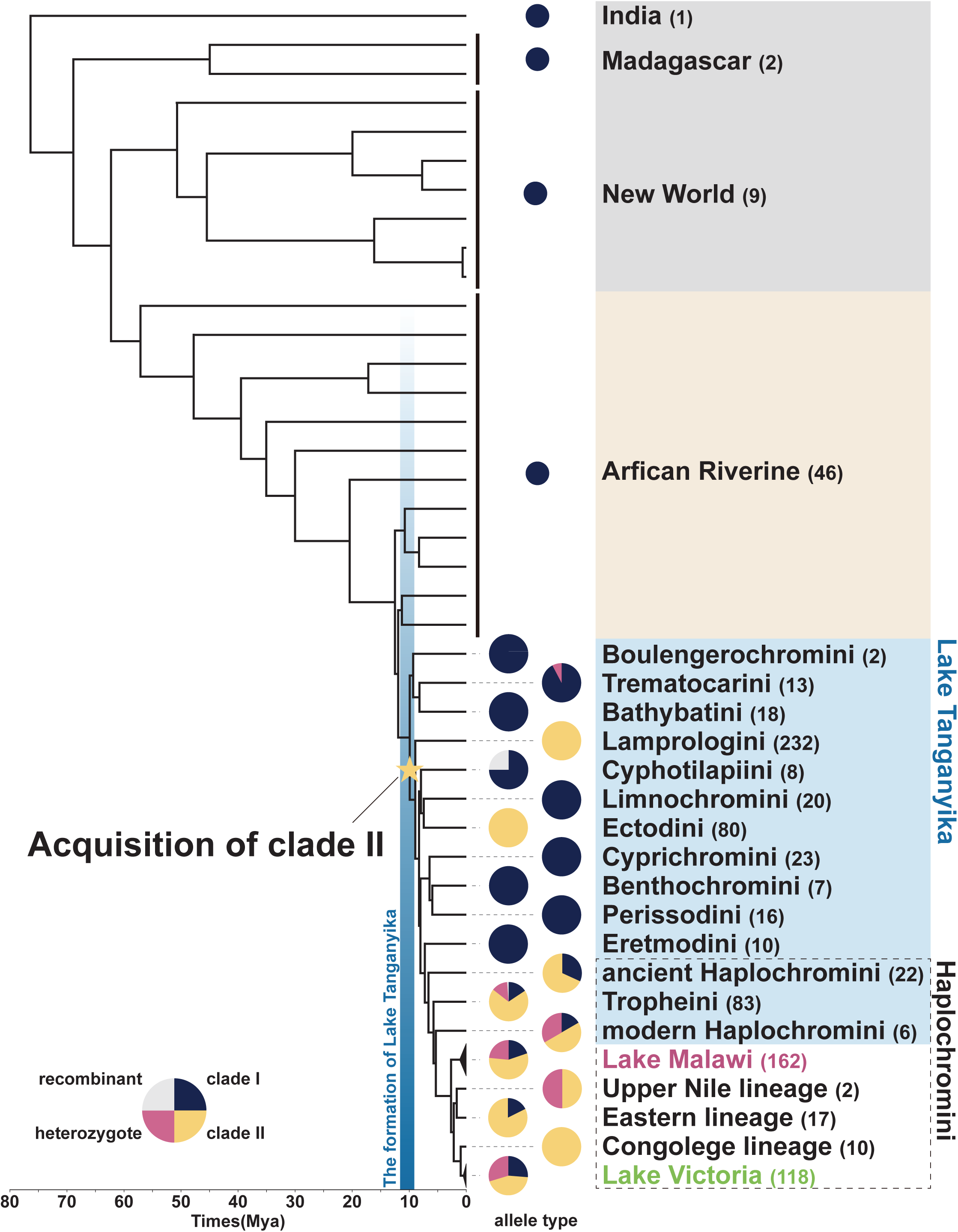
Frequencies of each allele of V1R2 in each cichlid lineage. We calculated the frequencies of Clade I and II alleles in each lineage. Clade I alleles are shown in dark blue, Clade II in yellow, heterozygotes in pink, and recombinants in gray. The phylogenetic tree and divergence times are based on previous studies (Matschiner et al., 2020; Meier et al., 2017; Nakamura et al., 2023; Ronco et al., 2021). Numbers in parentheses next to the lineage names represent the number of samples. The star represents the estimated acquisition times for Clade II.

### Highly variable alleles of V1R6 with multiple recombination in Lake Victoria cichlids

We constructed a V1R6 phylogenetic tree, and its topology was consistent with that of the species tree (Figure 3A). We found the Lake Victoria cichlids to exhibit a notable number of alleles, which were differentiated by a combination of 13 variant sites (Figure 3B). These allelic variants were present as polymorphisms across different species. Substitutions occurring independently multiple times at the same site for each allele are extremely unlikely. Therefore, we hypothesized that this wide range of alleles originated from recombination between two highly divergent alleles present in the ancestral population of Lake Victoria (Figure 3C). Accordingly, we found that the recombination rate of the genomic region surrounding V1R6, estimated using data from a previous study (Feulner et al., 2018), was higher than that of regions surrounding other V1R genes (Figure S8). Based on our hypothesis, we focused on the 13 variant sites identified in Lake Victoria cichlids and subsequently classified the V1R6 alleles of cichlids present across all the East African Great Lakes and rivers. We did not find any potential signature of recombination between highly divergent alleles in cichlids residing outside the Lake Victoria basin. Subsequently, we reconstructed the ancestral sequence at the node denoted by the arrowhead in Figure 3A and designated this sequence as the “ancestral allele” (Figure 3B, left top; dark blue nucleotide). We then subtracted the nucleotides of the “ancestral allele” (dark blue) from the 13 polymorphic sites of Lake Victoria cichlids to reconstruct a hypothetical “derived allele” (yellow), representing a counterpart sequence of the past recombination event (Figure 3B, right top). Accordingly, the alleles found in Lake Victoria cichlids were categorized as “ancestral,” “derived,” and their “mixtures.” Notably, cichlids from the Upper Nile, Eastern lineage, and Congolese lineage possess recombinant alleles similar to those of Lake Victoria cichlids (Figure 3B). However, despite extensive investigations, the hypothetical “derived allele” was not found in any extant cichlid, except for those in Lake Victoria (e.g., haplotypes Lake Victoria 51 and 52; Figure 3B, right bottom).

**Figure 3.**
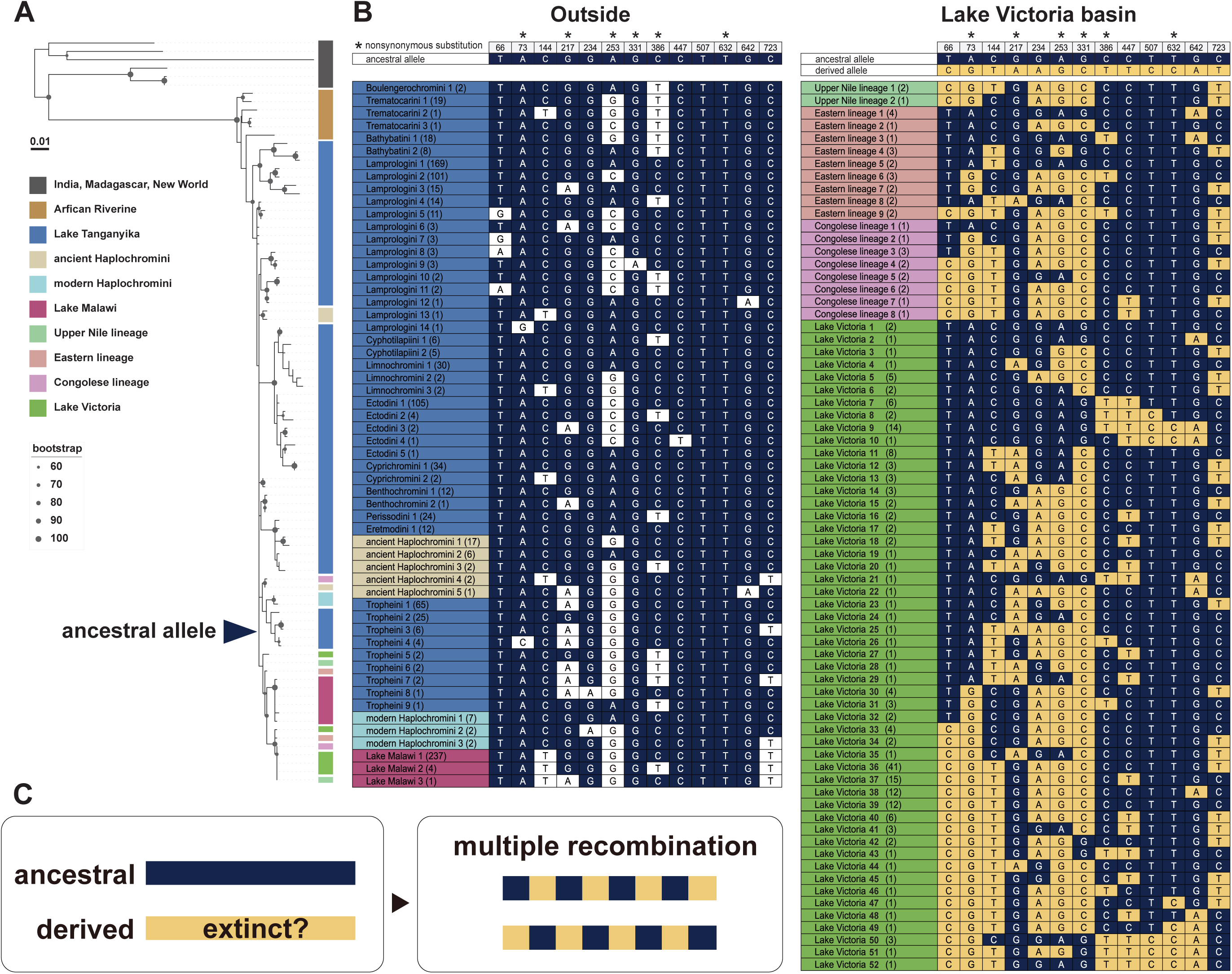
Recombination events observed in the V1R6 gene of cichlids. (A) Phylogenetic tree of the V1R6 gene. Bootstrap values are shown only for nodes with support values of ≥60. (B) All haplotypes of the V1R6 gene. Only sites that are polymorphic in Lake Victoria lineage are displayed. Numbers in parentheses following lineage names indicate the number of samples. Asterisks above the site numbers denote nonsynonymous substitutions. Ancestral and derived alleles at each site are defined based on the frequency of nucleotides across the sequences. Sites matching the ancestral allele are highlighted in dark blue, and those matching the derived allele are highlighted in yellow. (C) Schematic diagram of the recombination.

### Highly divergent alleles of V1R1, V1R3, and V1R6 in cichlids of particular tribes

We identified two highly divergent Tropheini V1R1 alleles, namely Clades I and II, which are distinguished based on 11 amino acids (Figure 4A, see Method). The monophyly of Clades I and II was corroborated by a maximal bootstrap probability (Figure S9). The amino acid sequence of Tropheini Clade I is similar to that of other cichlids; thus, Clades I and II are defined as “ancestral” and “derived” alleles, respectively. Furthermore, the dN/dS ratio of the branch where Clade II diverged from Clade I was significantly higher than that of the background branches (Table S2), indicating a positive selection acting on the divergent branch. We then calculated the frequencies of the two major V1R1alleles (Clades I and II) in Tropheini and found that Clades I and II were present in 28% and 65% of the Tropheini samples, respectively (Figure 4B). Extensive investigations of Clade II sequences revealed their presence only in Tropheini cichlids among the various cichlids examined. The polymorphic sites differentiating Clade I from Clade II in cichlids were highly conserved in Clade I of ray-finned fish (Figure S10).

**Figure 4.**
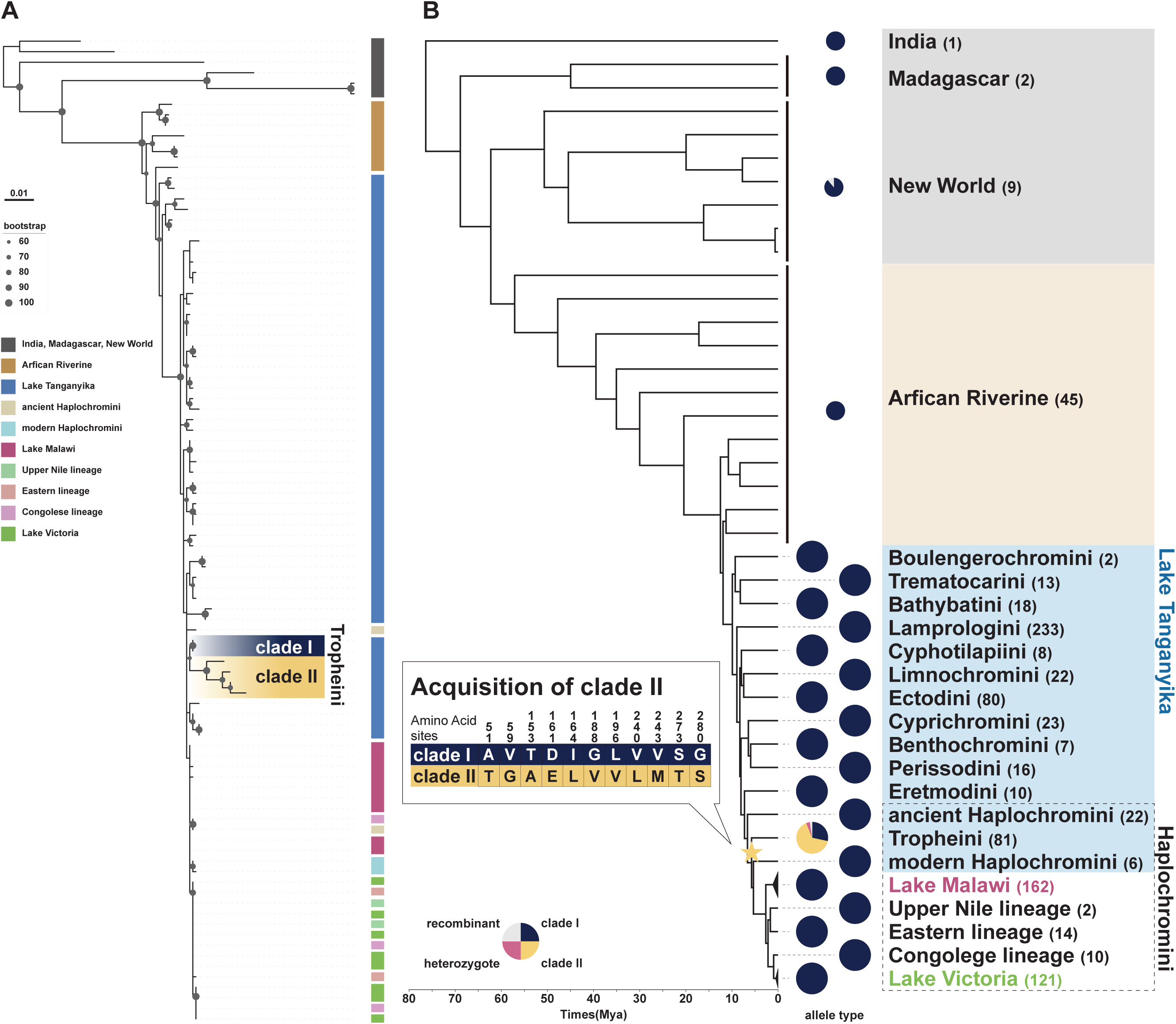
Two highly divergent alleles of V1R1 observed in Tropheini. (A) A phylogenetic tree of the V1R1 gene. Bootstrap values are shown only for nodes with support values of ≥60. (B) Frequencies of each allele of the V1R1 gene for each cichlid lineage. Clade I alleles are shown in dark blue, Clade II alleles in yellow, heterozygotes in pink, and recombinants in gray. The phylogenetic tree and divergence times are based on previous studies (Matschiner et al., 2020; Meier et al., 2017; Nakamura et al., 2023; Ronco et al., 2021). Numbers in parentheses next to the lineage names represent the number of samples. The star represents the estimated acquistion timing for Clade II.

We also identified highly divergent alleles of V1R3 in the Ectodini and Trematocarini. Seven nonsynonymous substitutions were identified on the branch corresponding to the common ancestor of Ectodini (Figure S11). This branch exhibited a significantly higher dN/dS ratio than the background branches (Table S2). Furthermore, 10 nonsynonymous substitutions were found on the branch representing the common ancestor of Trematocarini, excluding *Trematocara unimaculatum* (Figure S12). This branch exhibited a significantly higher dN/dS ratio than the background branches (Table S2). We found divergent alleles for V1R6 in the Ectodini and Trematocarini (Figure S13), with the dN/dS ratio of these clades (Ectodini, omega = 2.69; Trematocarini, omega = 1.61) being significantly higher than that of the background (Table S2).

### V1R2 amino acid substitution sites are located near the ligand-binding pocket

To understand the functional impact of the amino acid substitutions observed between the two major divergent alleles of V1R2, we modeled the 3D structure of the V1R2 protein and predicted its ligand-binding sites (Figure 5A). We first modeled the 3D structures using the amino acid sequences of Clades I and II and then used 4HPAA, a highly promising candidate ligand (Behrens et al., 2014; Kawamura & Nikaido, 2022), to determine the ligand-binding sites for each clade and compare them. Eight of the 11 amino acid substitution sites (TM2: C62→Ser, ECL1: Q65→Pro, TM4: A145→Ser and A146→Thr, ECL2: P159→Ser and H169→Leu, TM5: V185→Ile, and TM6: D247→Glu) we identified were situated near the predicted ligand pocket (Figure 5B). Furthermore, four of these sites (TM4: A145→Ser and A146→Thr, TM5: V185→Ile, and TM6: D247→Glu) overlapped with the ligand-binding sites (Figure 5B). Additionally, the amino acid substitutions of V1R2 were significantly skewed toward the ligand-binding sites (Fisher’s exact test: p = 0.0389).

**Figure 5.**
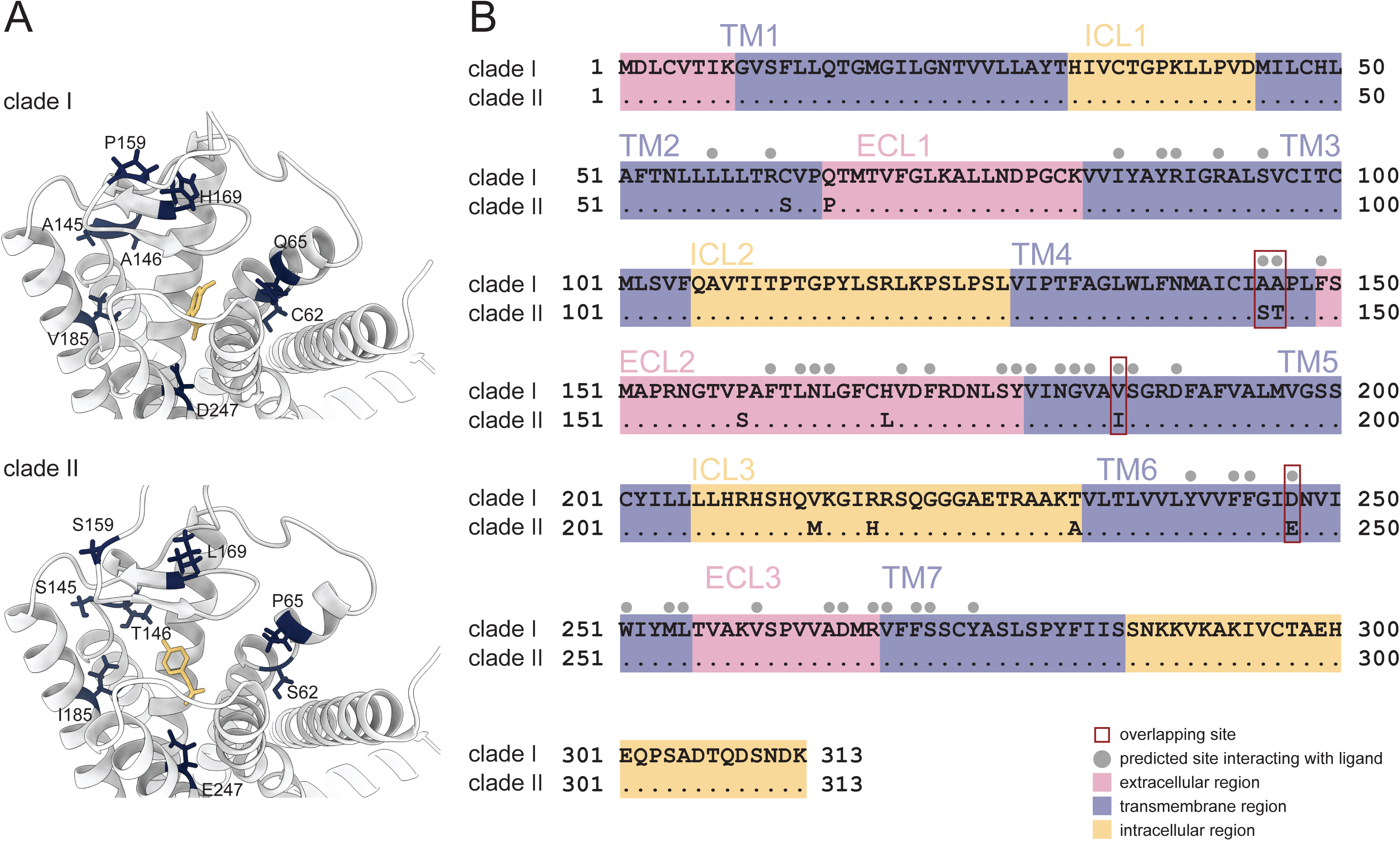
3D structure of cichlid V1R2. (A) 3D structure of V1R2 viewed from the extracellular side. Amino acids that differ between Clades I and II are shown in dark blue. The ligand used for analysis (4HPAA) is shown in yellow. (B) Predicted transmembrane regions of cichlid V1R2. The transmembrane regions were predicted using the Clade I sequence. Dots indicate amino acids identical to those in Clade I. The extracellular regions are shown in pink, transmembrane regions in blue, and intracellular regions in yellow. TM: transmembrane; ECL: extracellular loop; ICL: intracellular loop. Gray dots indicate the predicted ligand-binding sites. Red squares indicate ligand-binding sites that are also sites of substitution between alleles. Mutations occurred at ligand-binding sites at a significantly higher rate than that expected by chance (Fisher’s exact test: p = 0.0389).

We also performed the same analyses for V1R1 and V1R6 (Figure S14A, S15A). In V1R1, 5 of the 11 amino acid substitution sites (TM2: V59→Gly, ECL2: I164→Leu, TM5: G188→Val, and TM7: S273→Thr and G280→Ser) we identified were situated close to the predicted ligand pocket (Figure S14B). Furthermore, three of these sites (ECL2: I164→Leu; TM7: S273 → Thr and G280 →Ser) overlapped with the ligand-binding sites (Figure S14B). However, the skewed distribution of the V1R1 amino acid substitutions toward the ligand-binding sites did not show statistical significance (Fisher’s exact test: p = 0.0922). In V1R6, none of the amino acid substitution sites were located close to the predicted ligand pocket (Figure S15B).

## Discussion

### Allelic and functional diversity of V1R in East African cichlids

We conducted large-scale analyses of V1R diversity in cichlids using WGS data from 528 species and 907 samples, which have recently been made available following large-scale comparative genomic studies (Malinsky et al., 2018; McGee et al., 2020; Ronco et al., 2021). As a result, we identified an accumulation of nonsynonymous substitutions and signatures of positive selection in V1R1, V1R2, V1R3, and V1R6 (Figure 6A). Notably, two highly divergent alleles of V1R1 and V1R2 were found in the Tropheini (Figure 4B) and those inhabiting the East African Great Lakes (Figure 2B), respectively. In addition, highly divergent alleles of V1R3 and V1R6 were identified in the Trematocarini and Ectodini (Figure S11–13). These results corroborate previous research involving small-scale species sampling efforts (Nikaido et al., 2014).

**Figure 6.**
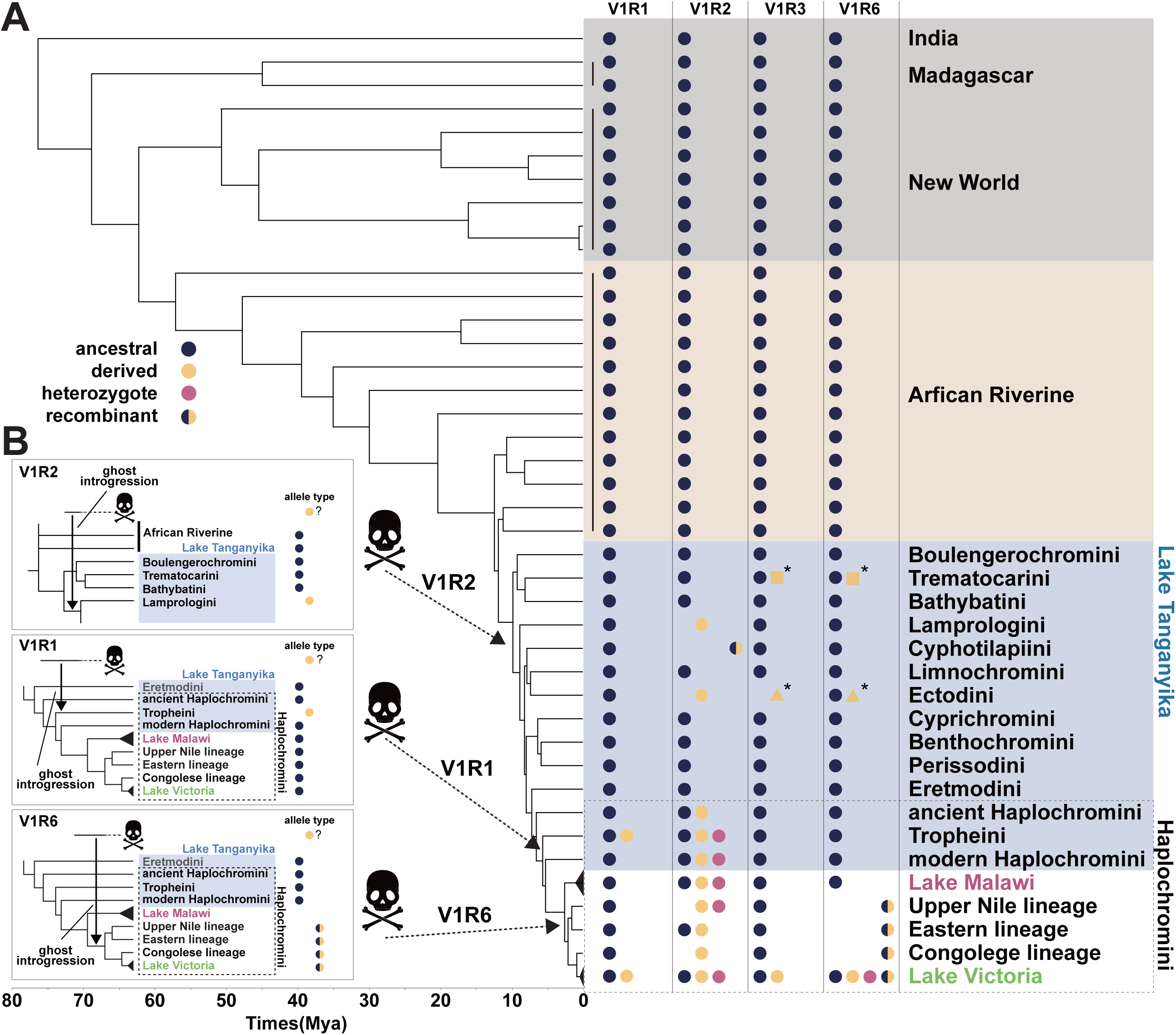
Possible involvement of ghost introgressions in cichlids (A) Sequence diversity of V1R genes in cichlids. The allele types of V1R genes are represented by colored dots for each lineage. The phylogenetic tree and divergence times are based on previous studies (Matschiner et al., 2020; Meier et al., 2017; Nakamura et al., 2023; Ronco et al., 2021). Note that the derived allele (Clade II) of V1R2 is shared across lineages, whereas the derived sequences of V1R3 and V1R6 differ among lineages (asterisks). (B) Predicted mechanisms of allele acquisition in the V1R1, V1R2, and V1R6 genes. Arrows indicate the most parsimonious estimated times of introgression.

The V1R gene orthologs are highly conserved across different fish species (Ota et al., 2012; Policarpo et al., 2022; Saraiva & Korsching, 2007; Zapilko & Korsching, 2016). The recently published V1R gene sequences of ray-finned fish (Policarpo et al., 2022) also exhibit high sequence similarity and low dN/dS ratios (Figure S16). Furthermore, the polymorphic sites that differentiate the divergent alleles of V1R1 and V1R2 in cichlids are highly conserved among ray-finned fish (Figure S7, S10). These results suggest that V1R gene diversification is unique to cichlids within the ray-finned fish category and is probably associated with rapid adaptive radiation.

The prediction of the 3D structure of V1R2 and the determination of the sites interacting with the ligand revealed that the amino acid residues differing between Clades I and II are located close to the ligand pocket. Furthermore, amino acid substitutions were predominantly found near the ligand-binding sites, showing a statistically significant trend (Figure 5B). Two amino acid substitutions were detected in ECL2 (extracellular loop 2) (Figure 5B), which is critical for ligand selectivity in class A G protein-coupled receptors (GPCRs), a category that encompasses V1R (Nicoli et al., 2022). ECL2 also contributes to ligand binding in the bitter taste receptor T2R, which is phylogenetically close to V1R (Xu et al., 2022). These findings suggest that the two divergent clades of V1R2 likely differ from one another in terms of their ligand-binding properties.

V1R6 exhibited the lowest sequence similarity and highest dN/dS ratio among the six V1R genes in all ray-finned fish species (Figure S16). Hence, the pressure of negative selection operating on V1R6 is weaker than that on other V1Rs, not only in cichlids but also across all ray-finned fish species. Notably, V1R6 of the Lake Victoria basin cichlids exhibited high variability, with numerous alleles resulting from multiple recombination events between divergent alleles (Figure 3B, 3C). The recombination rate in the V1R6 region of Lake Victoria cichlids was considerably high (Figure S8), indicating that V1R6 may be located in a recombination hotspot. In contrast, recombination in V1R1 and V1R2 is strongly suppressed. The structural constraint on V1R6 appears relatively weak, implying that the ligand specificity of V1R6 may be lower than that of other V1Rs. In the 3D structural analysis, the ligand pockets of V1R1 and V1R2 were predicted to be within the TM region, which is analogous to that found in typical class A GPCRs (Figure 5B, S14B). Conversely, the ECL2 region was predicted to facilitate ligand binding in V1R6 (Figure S15B). This suggests that V1R6 is distinct from V1R1 and V1R2 in terms of its functions and ligand binding mechanisms. Investigating the distinct structural differences predicted among V1Rs will provide insight into the function of V1Rs in fish, a topic of ongoing debate (Behrens et al., 2014; Cong et al., 2019; Saraiva & Korsching, 2007).

### The origin of striking V1R diversity through ghost introgression

Understanding the remarkable diversity of V1Rs in cichlids is crucial, particularly their origin and maintenance over the 10 million years of their evolutionary history. We first investigated the origin and process of V1R2 diversification by extensively investigating the existence of Clade I and II alleles in each cichlid lineage (Figure 2). Clade II was found in cichlid lineages inhabiting the East African Great Lakes but not in lineages that diverged prior to the Lake Tanganyika radiation. Specifically, the Lamprologini and Ectodini were fixed into Clade II, whereas those of other Lake Tanganyika tribes were fixed into Clade I. The cichlids of Haplochromini were homozygotes of Clade I or II, or heterozygotes of both. The pattern of V1R2 allele distribution indicates that Clade II emerged suddenly approximately 10 million years ago, coinciding with the radiation of Lake Tanganyika cichlids. Given that the adaptive radiation of cichlids in Lake Tanganyika was rapid, it is unlikely for 11 nonsynonymous substitutions to accumulate gradually in a single allele within this timeframe, even under the influence of positive selection. Therefore, it is parsimonious to assume that the Clade II allele was already present in some populations/species before the radiation event and was subsequently introgressed into Lake Tanganyika cichlids through hybridization.

The genomes of East African cichlids contain several examples of standing genetic variation (SGV) (Nakamura et al., 2021; Svardal et al., 2021), including some that date back to before the adaptive radiation events in Lake Tanganyika (Nakamura et al., 2023). Clade I and II alleles of V1R2 represent clear examples of this old SGV. One of the most plausible explanations for the origin of this SGV is the melting-pot hypothesis (Nakamura et al., 2023; Weiss et al., 2015). According to this hypothesis, several cichlid populations/species, previously scattered throughout wetlands and small lakes, hybridized just before the adaptive radiation event in Lake Tanganyika. If each scattered population became fixed for Clades I and II, their hybridization may have provided the origin of the SGV for the highly divergent V1R2 alleles. However, one of the two alleles comprising the SGV of V1R2, Clade II, was not detected despite an extensive investigation of the 46 basal lineages of the riverine species (Figure 2). The absence of the Clade II allele in the riverine species suggests that the transporter lineage of the Clade II allele is already extinct.

Gene flow from extinct lineages, called ghost introgression, has been identified in various vertebrates, from gobiid fish to elephants, in recent years (Ai et al., 2015; Frei et al., 2022; Gopalakrishnan et al., 2018; Kato et al., 2023; Palkopoulou et al., 2018; Zhang et al., 2019). The sudden emergence of Clade II of V1R2 in Lake Tanganyika cichlids can be attributed to ghost introgression. Introgressed alleles are usually identified as highly divergent from those in nonintrogressed genomic regions. However, neutral genomic regions frequently undergo recombination and genetic drift following introgression, thereby complicating the identification of introgression signatures. Accordingly, V1R2 serves as a good example for identifying introgression due to its strict constraints on recombination. African Riverine lineages that predate adaptive radiation are referred to as “transporters” of genetic diversity (Loh et al., 2013). Notably, gene flow from Steatocranini, a riverine lineage that diverged before adaptive radiation (Irisarri et al., 2018), coincided with the emergence of Clade II of V1R2. Therefore, the high diversity of V1R2 alleles (Clade II) is likely a result of an introgression event from an extinct African Riverine lineage that diverged before adaptive radiation (ghost introgression) or from unexamined lineages (Figure 6B).

Our findings based on V1R6 data suggest that a significant number of distinct alleles emerged from the repeated recombination of 2 highly divergent alleles, one ancestral and the other derived, in the cichlids of the Lake Victoria basin, which were differed by 13 nucleotide substitutions (Figure 3B). Recombination of the LWS opsin gene was observed in Lake Victoria cichlids, which was attributed to hybridization between the Congo and Upper Nile lineages (Meier et al., 2017). In contrast, since both the Congo and Upper Nile lineages exhibit recombinant sequences, recombination between the ancestral and derived alleles of V1R6 must have occurred earlier than the hybridization event of the Congo and Upper Nile lineages. However, all the extant Haplochromini species, except Lake Victoria cichlids, lack the derived allele. Since the estimated mutation rate in cichlids is 3.5×10^−9^ (per site per generation), it is unrealistic for 13 substitutions to occur within the short timeframe of the emergence of the cichlids of Lake Victoria basin (∼400,000 years ago) (Johnson et al., 1996; Salzburger et al., 2014). Therefore, the derived allele of V1R6 was likely to have been introgressed from an extinct lineage (Figure 6B). The highly divergent alleles of V1R1 in Tropheini, as well as those of V1R3 and V1R6 in Ectodini and Trematocarini, are probably due to ghost introgression, similar to the cases of V1R2 and V1R6.

Notably, the nucleotide sequence differences between the two divergent alleles of V1R1, V1R2, and V1R6 were mostly >1% (V1R1 = 1.14%, V1R2 = 1.17%, and V1R6 = 1.39%). Therefore, each divergent allele may have been introduced into the ancestral population through a single ghost introgression event (melting-pot), following which the derived alleles would have independently sorted into separate lineages. Alternatively, introgression events in each lineage may have introduced the derived alleles of V1R1, V1R2, and V1R6 (Figure 6A, 6B).

## Conclusion

This study involved a comprehensive analysis of the V1R genes in East African cichlid species spanning numerous lineages, and we identified large-scale SGVs that can be attributed to ghost introgression. However, further research involving genome-wide statistical analyses rather than gene-level investigations of ghost introgression and melting-pot scenarios will be required to validate our findings. Our findings also highlighted the differences in structural constraints and selection pressure among the different V1Rs in cichlids, suggesting a functional distinction. This study provides important insights into the potential for V1R-mediated speciation and the genetic basis for adaptive evolution in cichlids.

## Acknowledgments

This study was supported by JSPS KAKENHI (20KK0167, 24K02074, 24KK0141) to M.N. Phylogenetic analyses were partially performed using the NIG supercomputer at ROIS National Institute of Genetics.

## Author Contributions

M.N. supervised the project. S.T. conducted all bioinformatics and evolutionary analyses. S.T. and M.N. wrote the manuscript. Z.Z., M.A., and T.N. provided their expertise and commented on the paper. All authors read and approved the final manuscript.

## Conflict of Interest Statement

The authors declare no conflict of interest.

## Data Availability Statement

V1R gene sequences of cichlids are available on Dryad (DOI). Scripts for obtaining V1R gene sequences of cichlids are available on Github (https://github.com/taki-sh/RegionCall).

---

**Figure S1.**
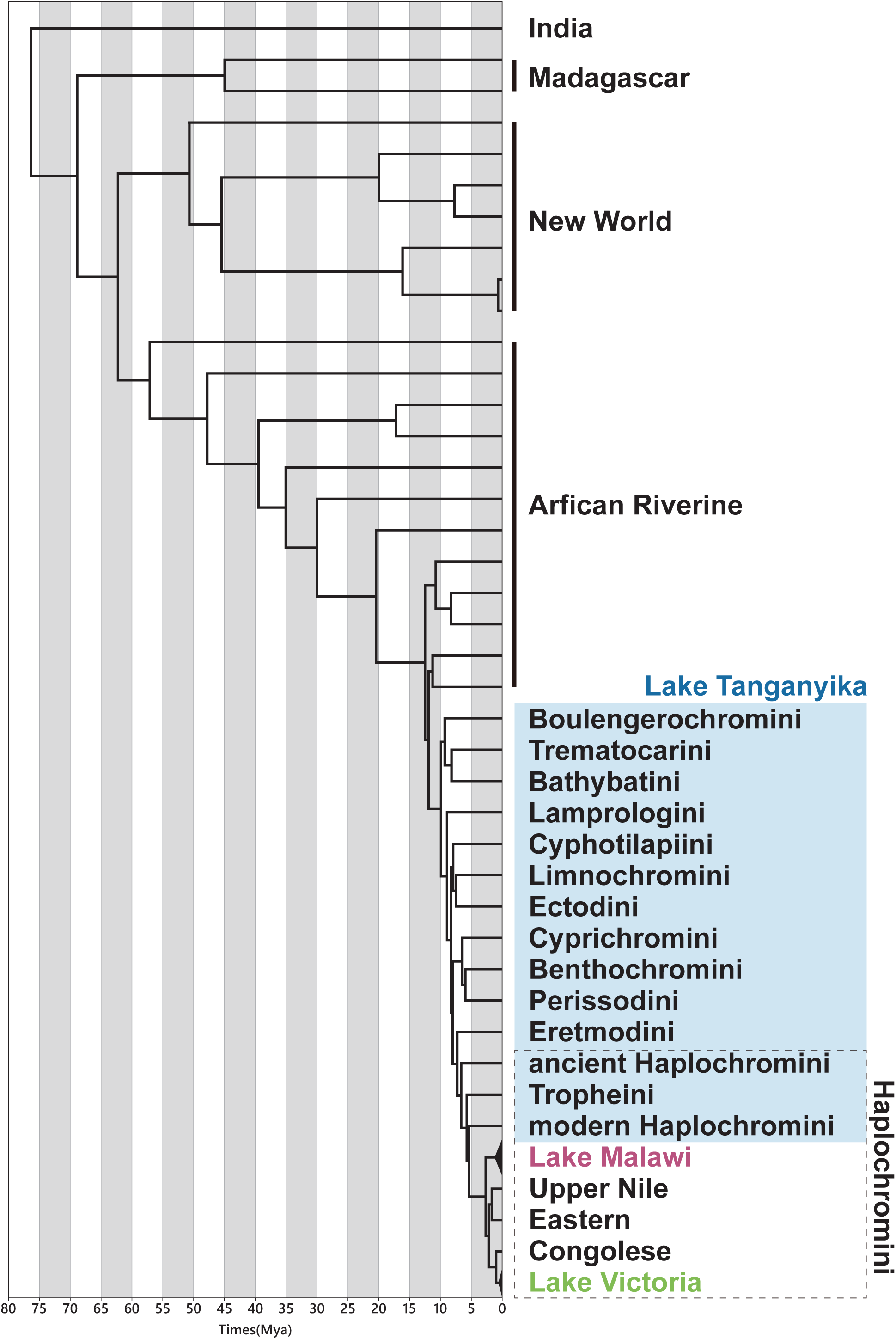
Lineage names in this study are defined with reference to previous studies (Matschiner et al., 2020; Meier et al., 2017; Nakamura et al., 2023; Ronco et al., 2021). Haplochromini species that diverged before Tropheini are defined as “ancient Haplochromini.” Divergence times are expressed in millions of years (Mya).

**Figure S2.**
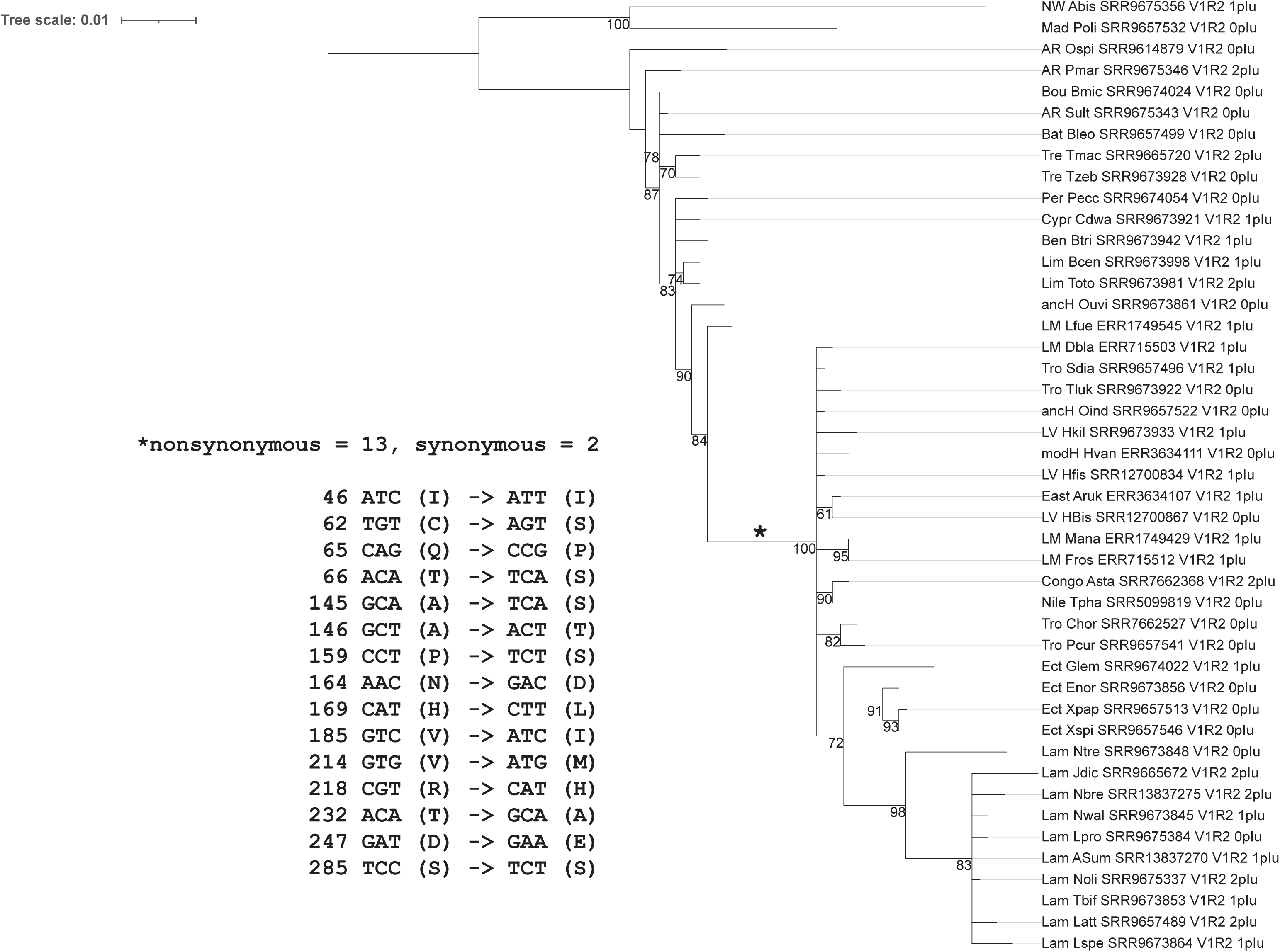
Phylogenetic tree and sequence alignment of the V1R2 gene in cichlids (Dataset 1). The phylogenetic tree displays bootstrap values only for nodes with support values of ≥60. The left panel shows nonsynonymous and synonymous substitution sites at positions marked with an asterisk.

**Figure S3.**
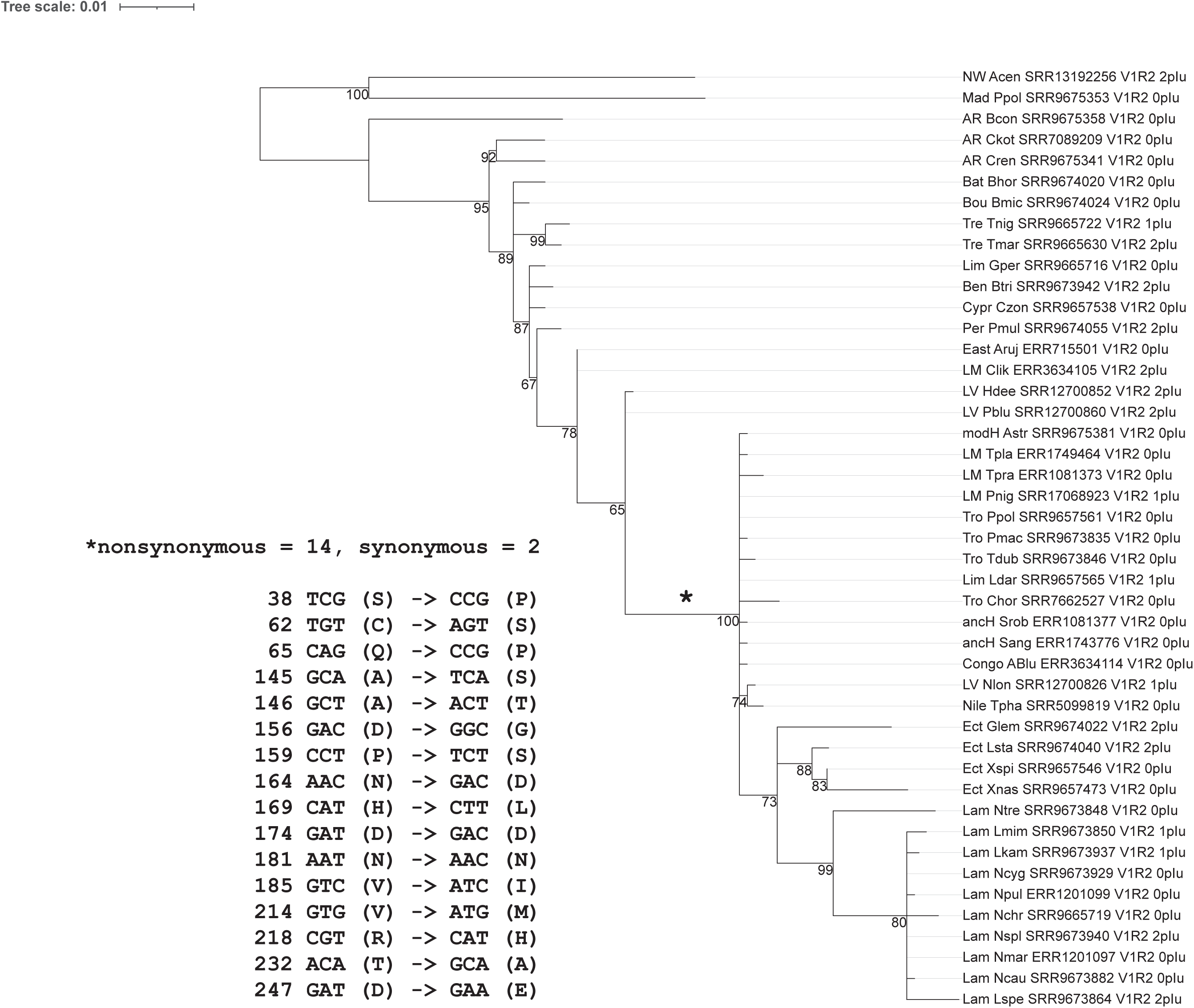
Phylogenetic tree and sequence alignment of the V1R2 gene in cichlids (Dataset 2). The phylogenetic tree displays bootstrap values only for nodes with support values of ≥60. The left panel shows nonsynonymous and synonymous substitution sites at positions marked with an asterisk.

**Figure S4.**
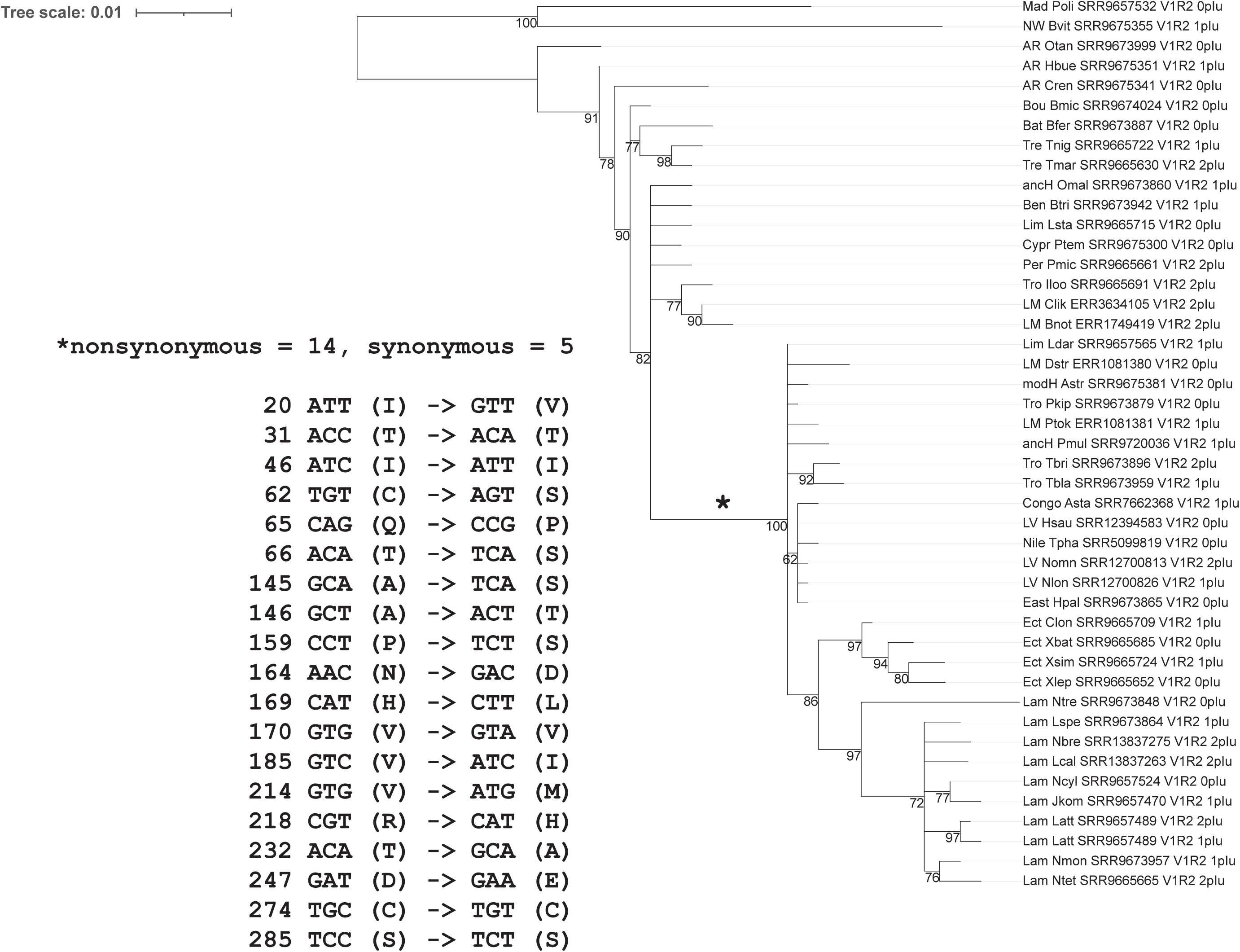
Phylogenetic tree and sequence alignment of the V1R2 gene in cichlids (Dataset 3). The phylogenetic tree displays bootstrap values only for nodes with support values of ≥60. The left panel shows nonsynonymous and synonymous substitution sites at positions marked with an asterisk.

**Figure S5.**
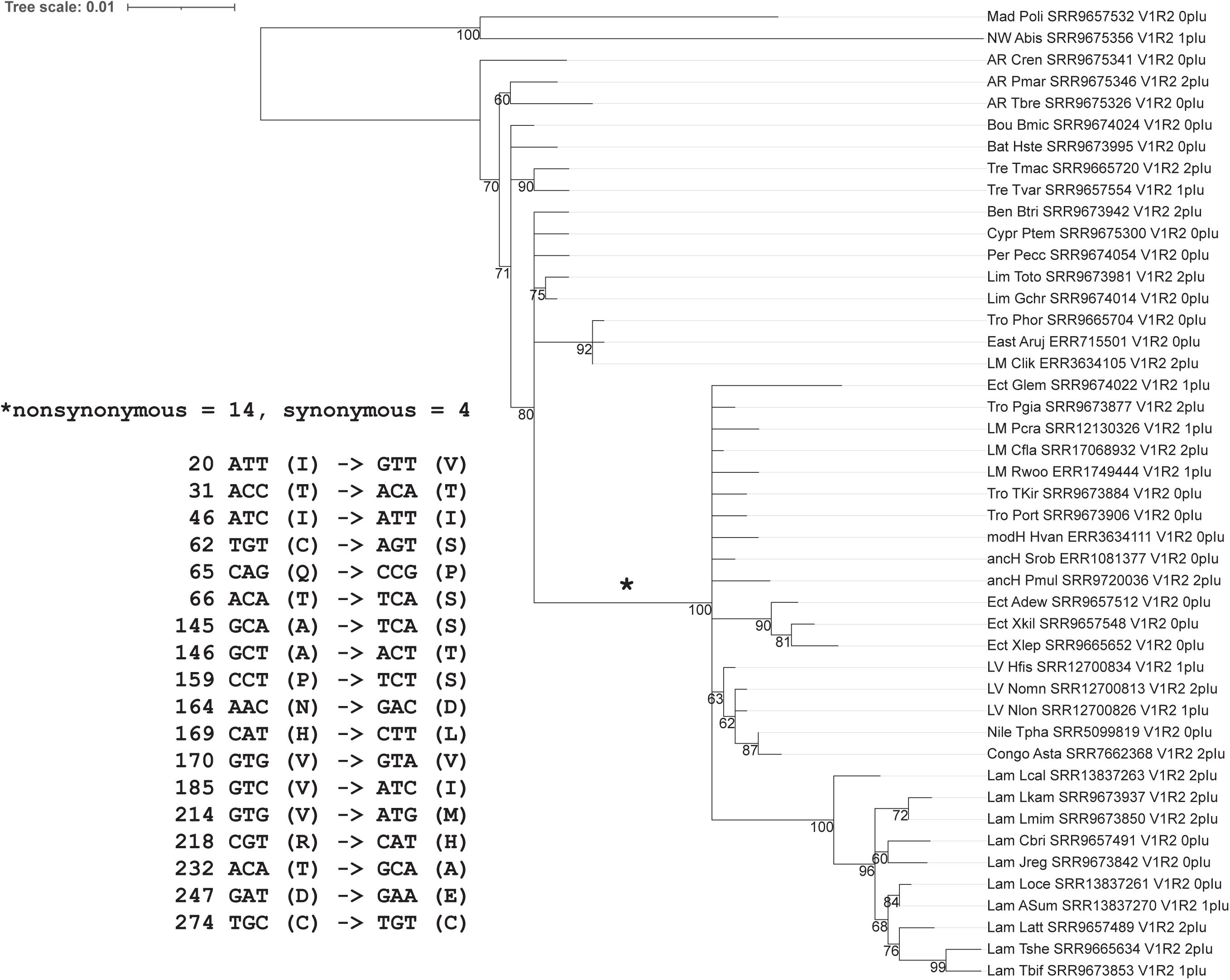
Phylogenetic tree and sequence alignment of the V1R2 gene in cichlids (Dataset 4). The phylogenetic tree displays bootstrap values only for nodes with support values of ≥60. The left panel shows nonsynonymous and synonymous substitution sites at positions marked with an asterisk.

**Figure S6.**
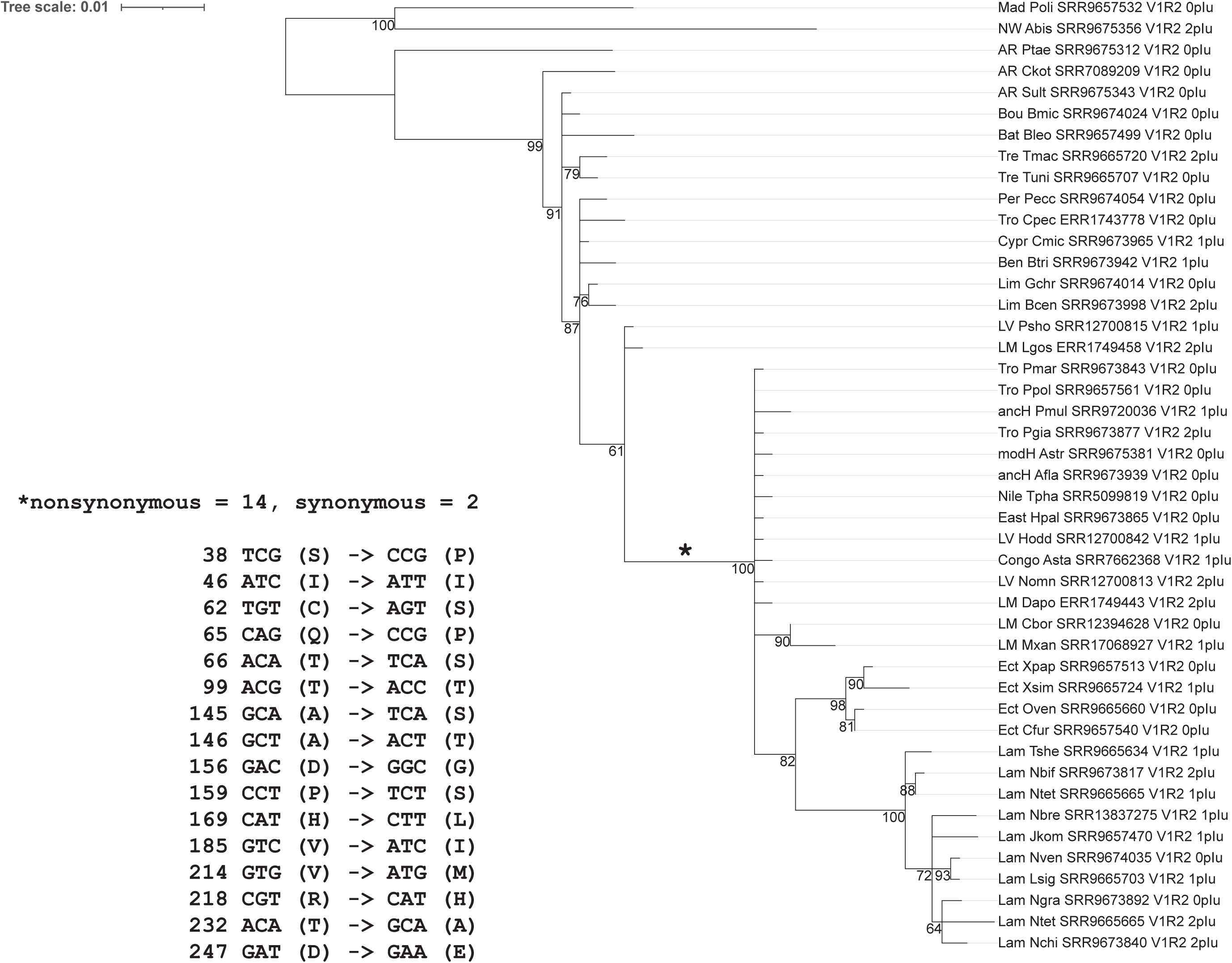
Phylogenetic tree and sequence alignment of the V1R2 gene in cichlids (Dataset 5). The phylogenetic tree displays bootstrap values only for nodes with support values of ≥60. The left panel shows nonsynonymous and synonymous substitution sites at positions marked with an asterisk.

**Figure S7.**
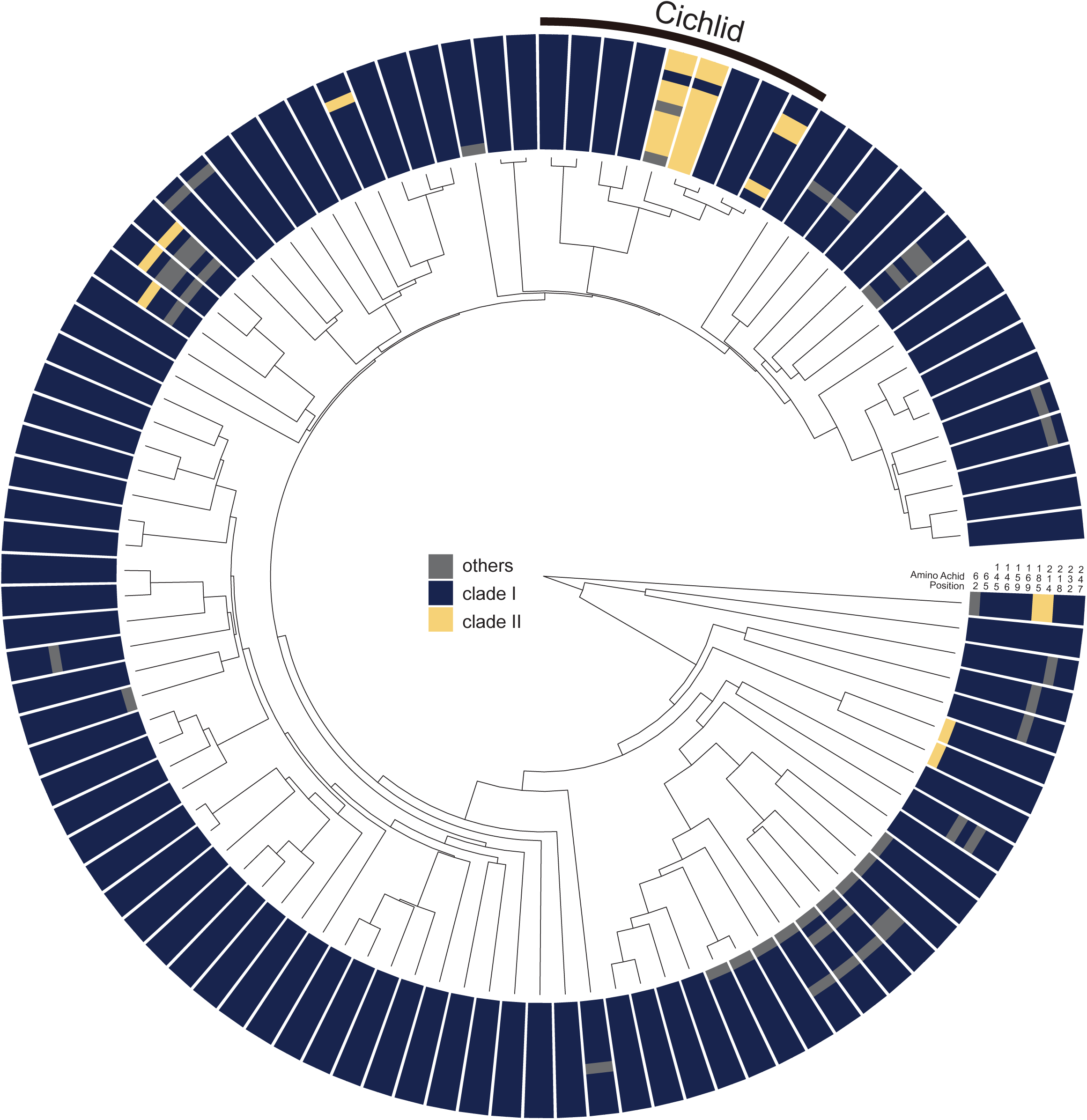
V1R2 alignment for 102 ray-finned fish species. The alignment displays sites corresponding to the two alleles (Clades I and II). Sites matching Clade I are highlighted in dark blue, those matching Clade II are highlighted in yellow, and all other sites are shown in gray.

**Figure S8.**
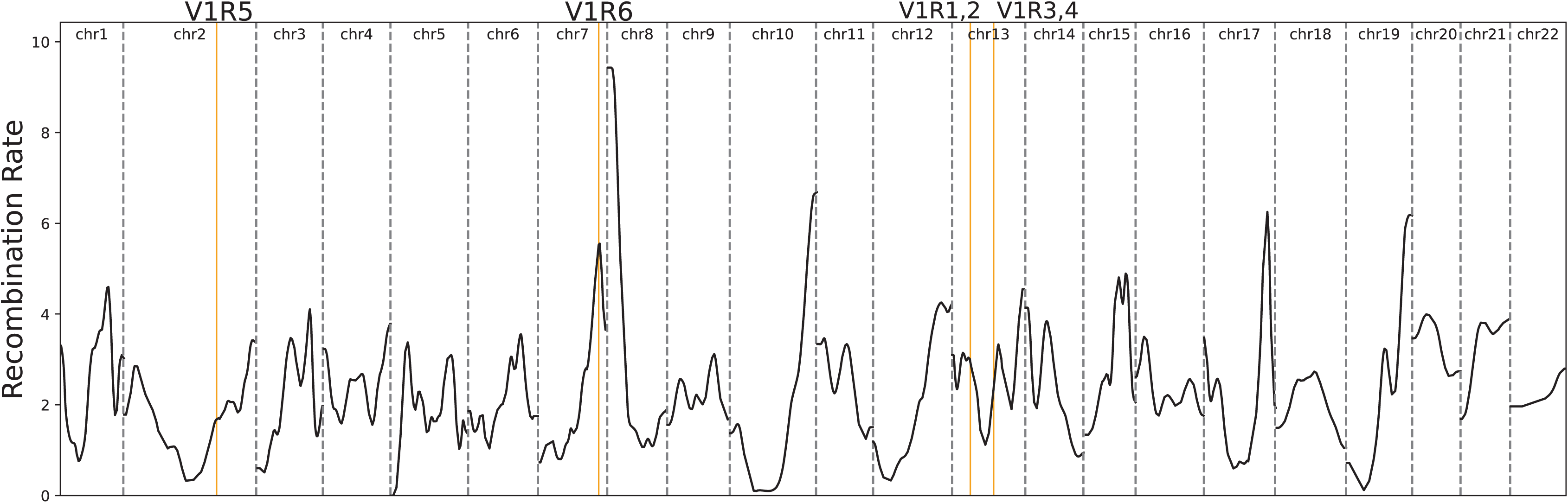
Recombination rate (cM/Mb) per chromosome in *Pundamilia nyererei* from the Lake Victoria lineage (Feulner et al., 2018). The positions of V1R genes are indicated by yellow lines.

**Figure S9.**
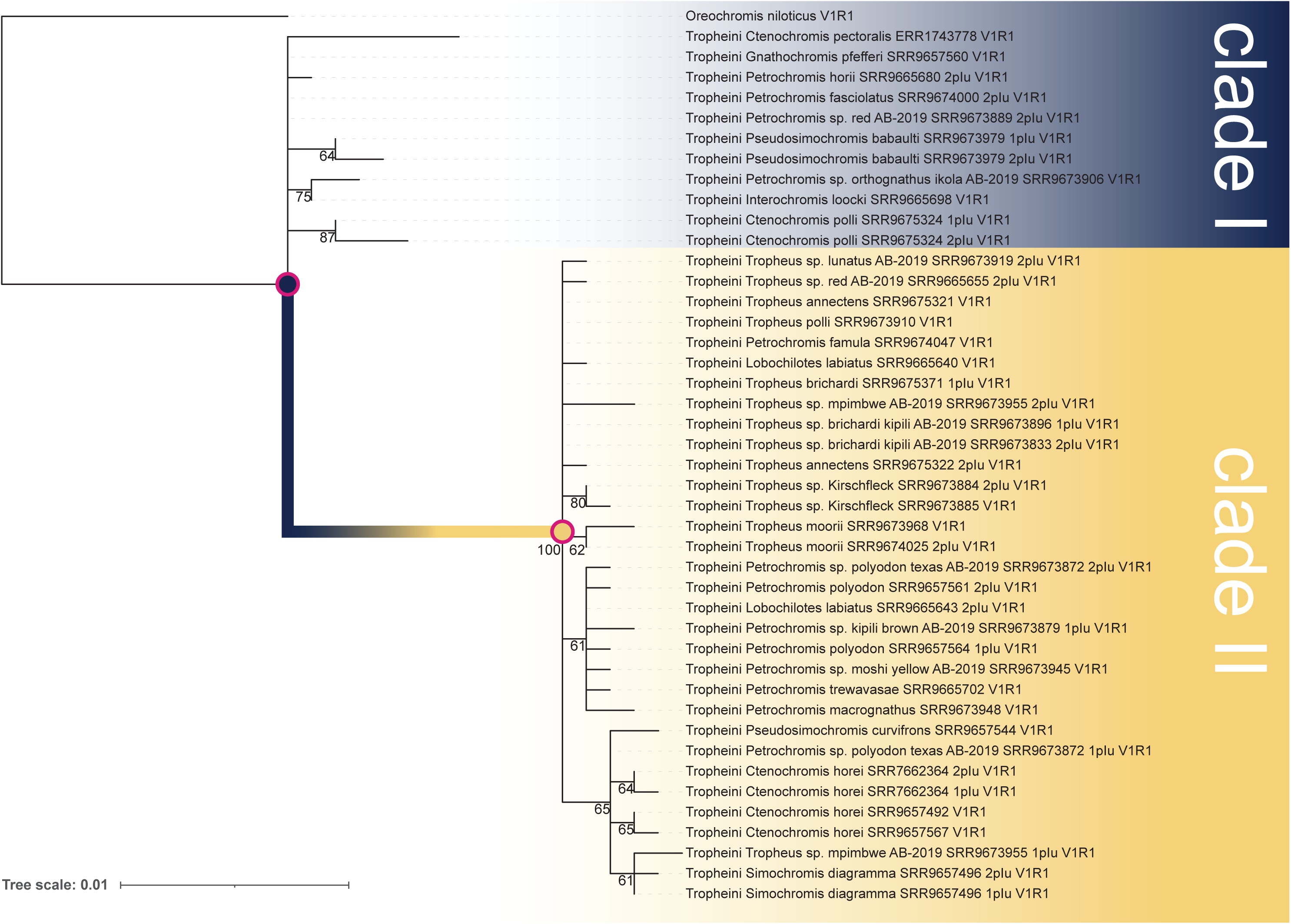
Phylogenetic tree of the V1R1 gene in Tropheini. Bootstrap values are shown only for nodes with support values of ≥60. The sequences are divided into two highly divergent alleles (Clades I and II). Recombination-derived sequences have been excluded.

**Figure S10.**
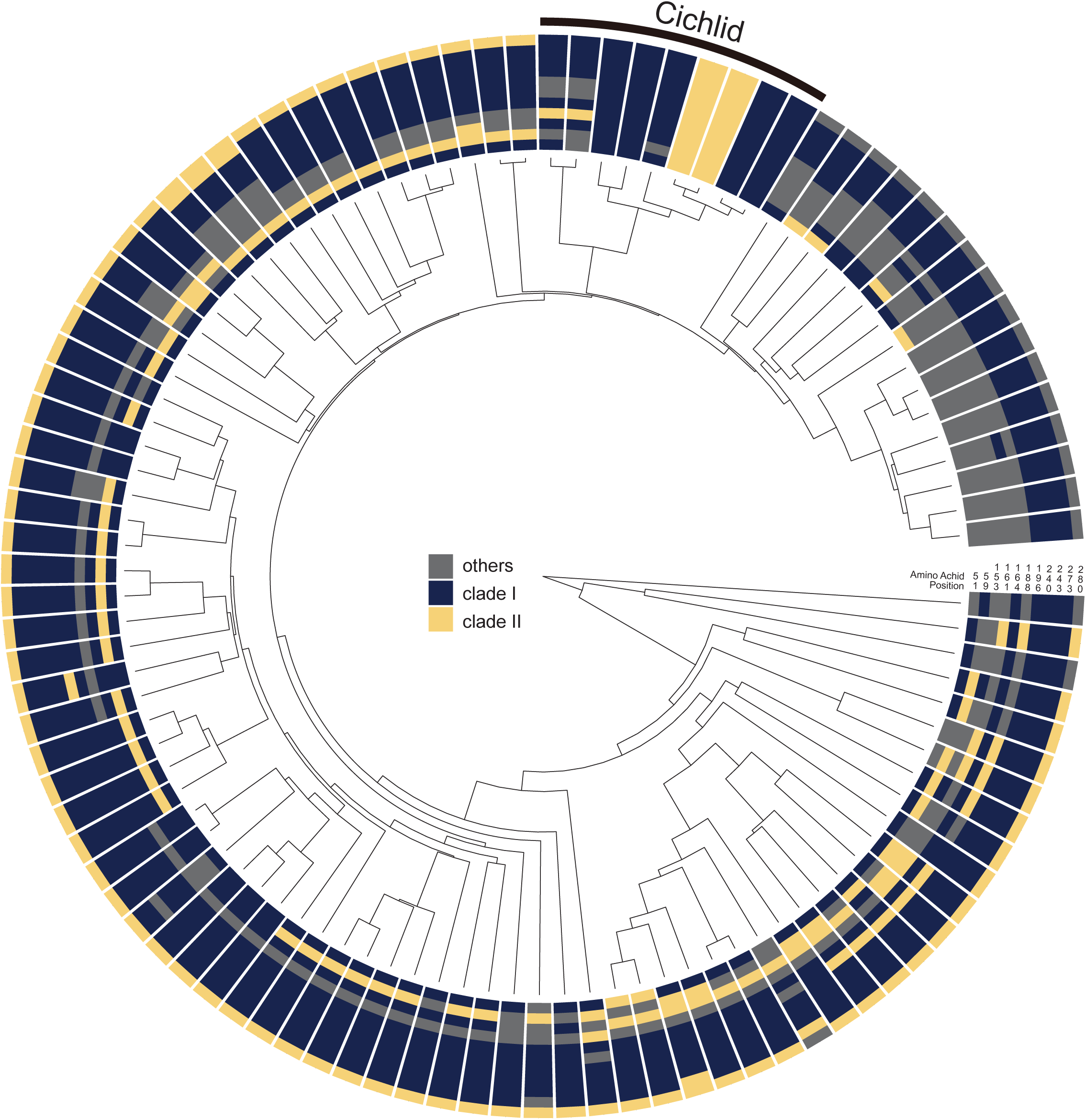
V1R1 alignment for 102 ray-finned fish species. The alignment displays sites corresponding to the two alleles (Clades I and II). Sites matching Clade I are highlighted in dark blue, those matching Clade II are highlighted in yellow, and all other sites are shown in gray.

**Figure S11.**
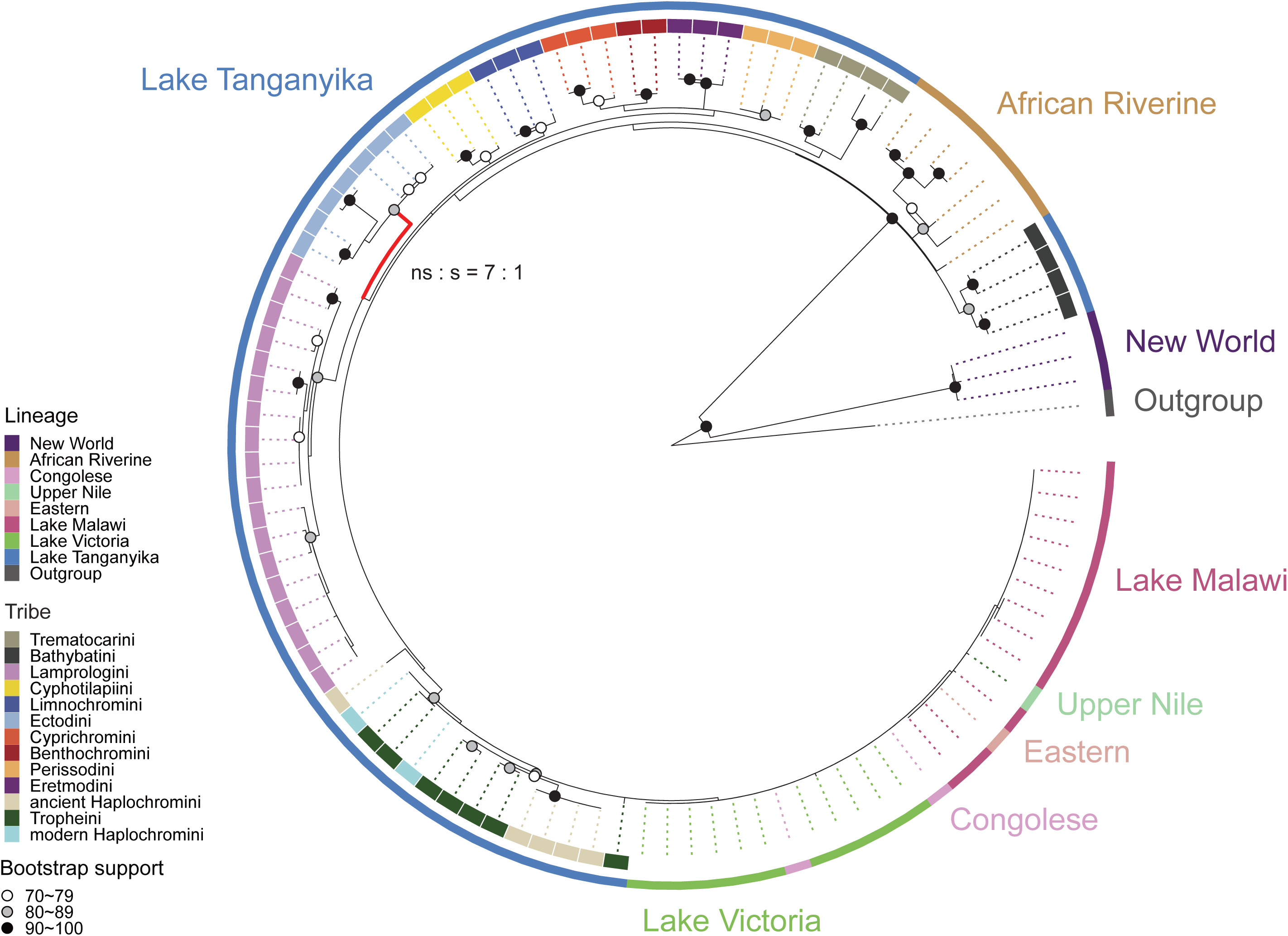
Phylogenetic tree of the V1R3 gene in cichlids. Nodes are marked with dots based on bootstrap values: black for values of ≥90, gray for 80–89, and white for 70–79. The inner circle represents tribes, whereas the outer circle indicates habitats. Branches under positive selection are highlighted in red (p = 0.0361, see Table S2). The number of nonsynonymous and synonymous substitutions for the branch is indicated as ns:s.

**Figure S12.**
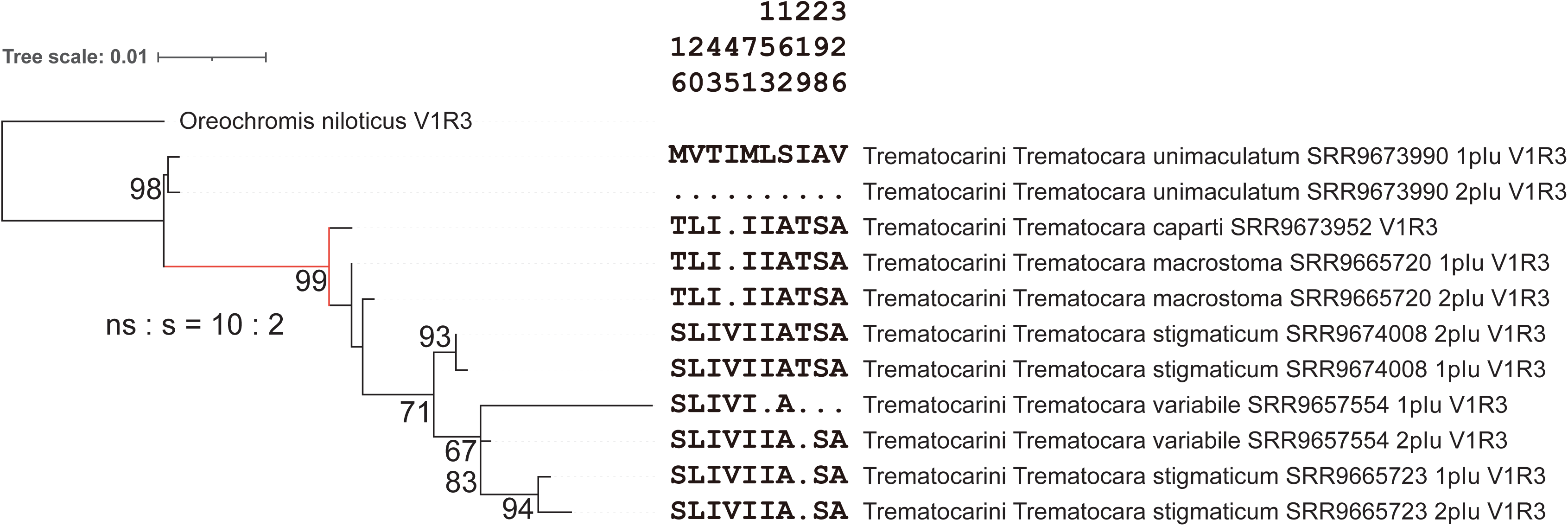
Phylogenetic tree and sequence alignment of the V1R3 gene in Trematocarini. Bootstrap values are shown only for nodes with support values of ≥60. *Trematocara unimaculatum*, which retains the ancestral sequence of V1R3, differs significantly from those of other species. The amino acid alignment displays only the sites with variations observed within Trematocarini. Dots indicate identity with the top sequence. Positive selection was not detected in the red branch (p = 0.0503, see Table S2). The number of nonsynonymous and synonymous substitutions in the red branch is indicated as ns:s.

**Figure S13.**
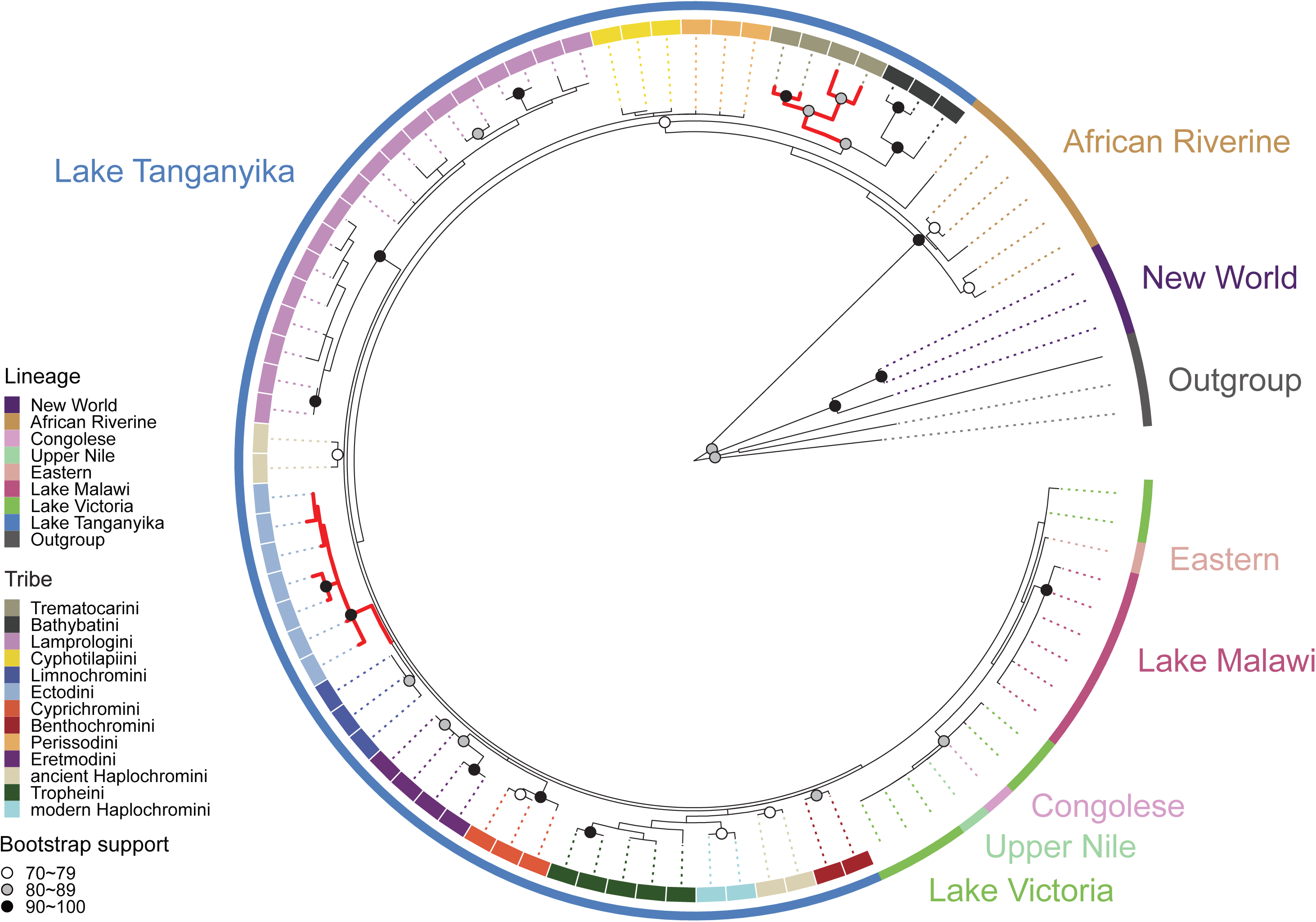
Phylogenetic tree of the V1R6 gene in cichlids. Nodes are marked with dots based on bootstrap values: black for values of ≥90, gray for 80–89, and white for 70–79. The inner circle represents tribes, whereas the outer circle indicates habitats. Branches under positive selection are highlighted in red (see Table S2).

**Figure S14.**
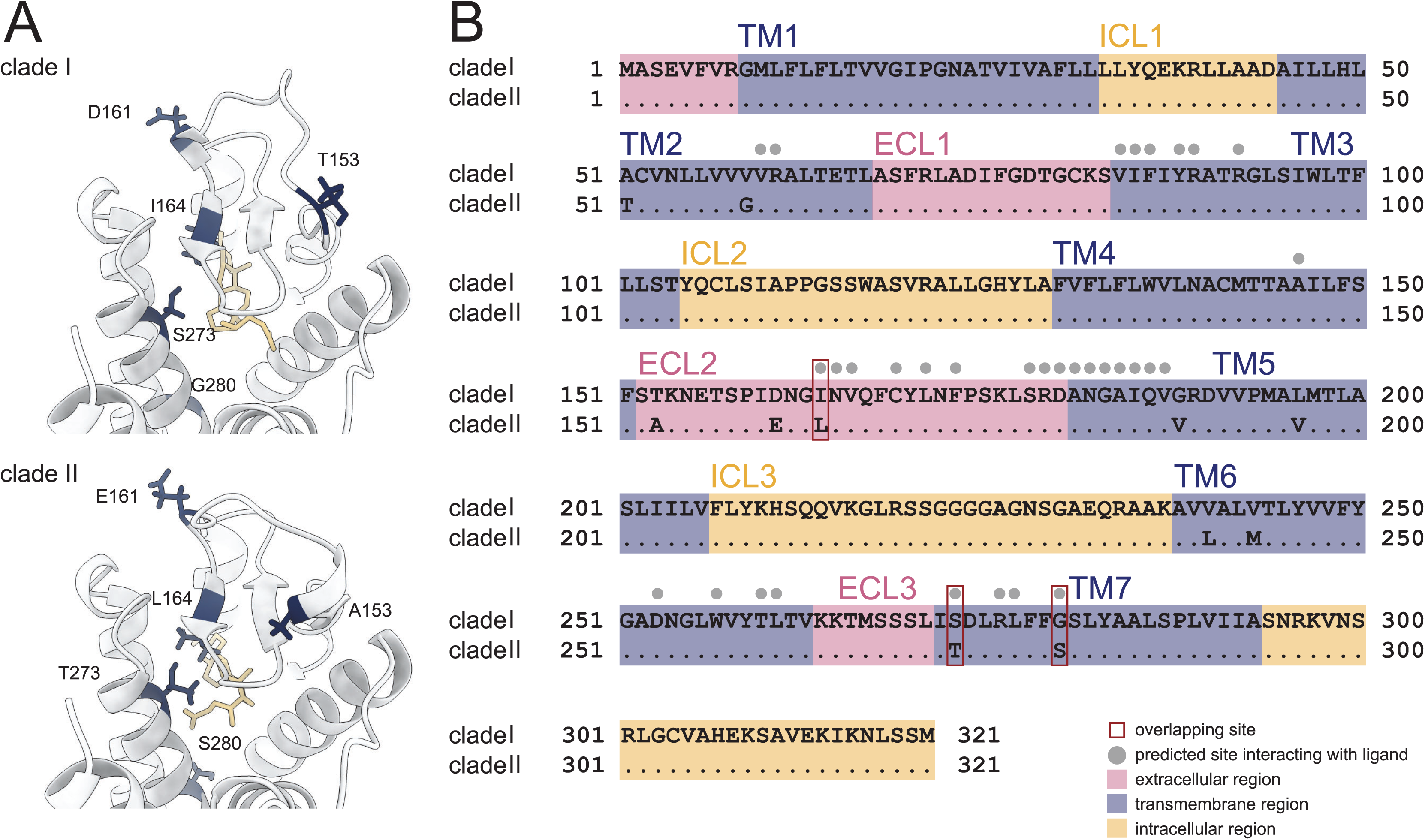
3D structure of cichlid V1R1. (A) 3D structure of V1R1 viewed from the extracellular side. Amino acids that differ between Clades I and II are shown in dark blue. The ligand used for analysis (lithocholic acid) is shown in yellow. (B) Predicted transmembrane regions of cichlid V1R1. The transmembrane regions were predicted using the Clade I sequence. Dots indicate amino acids identical to those in Clade I. The extracellular regions are shown in pink, transmembrane regions in blue, and intracellular regions in yellow. TM: transmembrane; ECL: extracellular loop; ICL: intracellular loop. Gray dots indicate the predicted ligand-binding sites. Red squares indicate ligand-binding sites that are also sites of substitution between alleles. (Fisher’s exact test: p = 0.0922)

**Figure S15.**
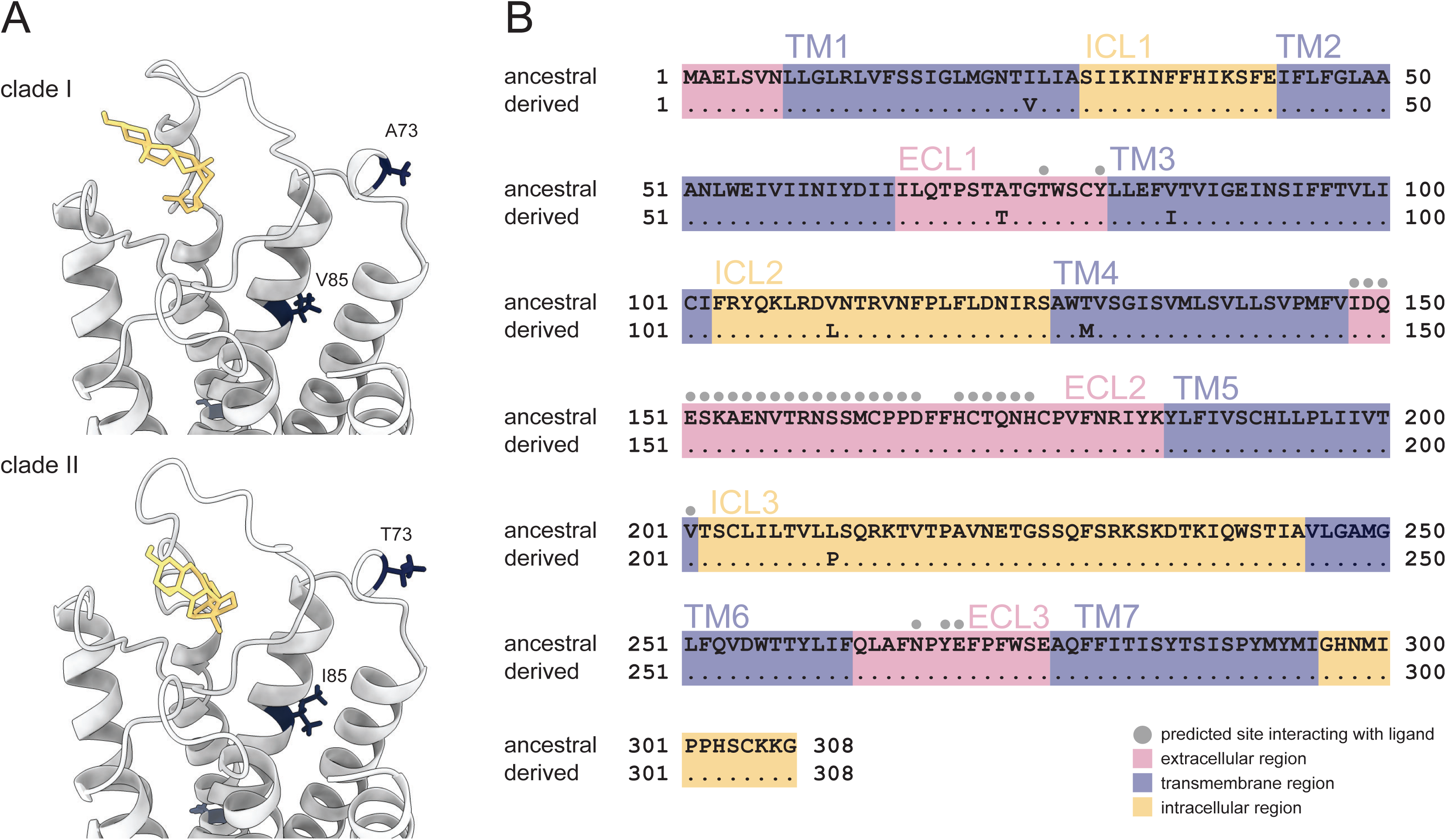
3D structure of cichlid V1R6. (A) 3D structure of V1R6 viewed from the extracellular side. Amino acids that differ between ancestral and derived alleles are shown in dark blue. The ligand used for analysis (lithocholic acid) is shown in yellow. (B) Predicted transmembrane regions of cichlid V1R6. The transmembrane regions were predicted using the ancestral allele sequence. Dots indicate amino acids identical to those in the ancestral allele. The extracellular regions are shown in pink, transmembrane regions in blue, and intracellular regions in yellow. TM: transmembrane; ECL: extracellular loop; ICL: intracellular loop. Gray dots indicate the predicted ligand-binding sites.

**Figure S16.**
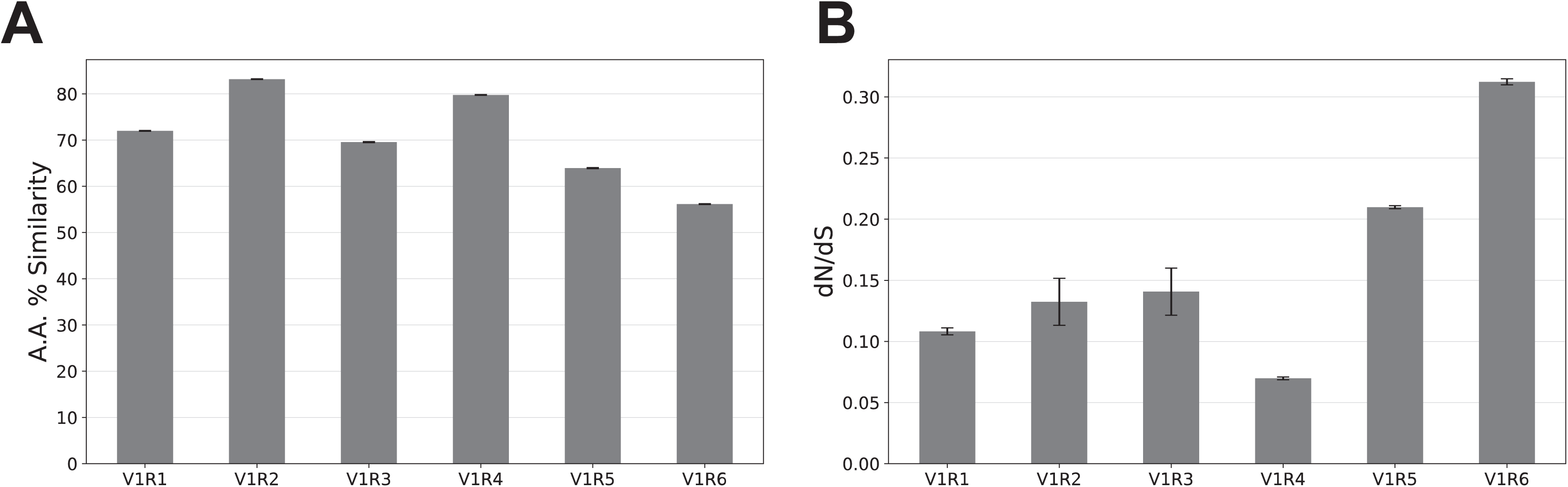
Sequence similarity and dN/dS ratios of V1R genes among 102 ray-finned fish species (A) Amino acid similarity of V1R genes. The average sequence similarity was calculated for all pairwise combinations among the 102 species. Error bars represent the standard errors. (B) dN/dS ratios of V1R genes. The average dN/dS ratio was calculated for all pairwise combinations among the 102 species. Error bars represent the standard errors.

Table S1 Short-read data from 28 tribes, 528 species, and 908 samples of cichlids registered in the NCBI Sequence Read Archive (SRA).

Table S2 Selection analyses and likelihood ratio test results.

## Notes

### Competing Interest Statement

The authors have declared no competing interest.

### Summary of Updates

The manuscript has been revised throughout for clarity and language, but the results remain unchanged

